# Integrative Genomic, Single-Cell, and Functional Profiling of the CD48–CD244 Axis and NK-Cell Dysfunction in Multiple Myeloma

**DOI:** 10.64898/2026.05.08.723909

**Authors:** Bonell Patiño-Escobar, Torsten Steinbrunn, Lucy Perez-Lugo, Sham Rampersaud, Daniel D. Waller, Huimin Geng, Fernando Salangsang, Paul Phojanakong, Juan Antonio Camara Serrano, Yan Zeng, Veronica Steri, Oscar A. Aguilar, Constantine S. Mitsiades, Arun P. Wiita

## Abstract

Multiple myeloma (MM) orchestrates immune evasion by subverting natural killer (NK) cell function. CD48, one of the most abundant NK-ligands on MM cells, paradoxically enhances NK-cell activation yet is associated with high-risk cytogenetics and poor patient survival. We integrated multi-omics (bulk and single-cell RNA-seq, ATAC-seq), genome-wide CRISPR-KO/a screens, and machine learning to dissect CD48 regulation and function. In human MM and Vκ*MYC mice scRNA-seq datasets, NK cells exhibit stepwise increases in inflammatory and exhaustion signatures and loss of cytotoxic potential as disease progresses. In vitro co-culture assays show CD48 overexpression on MM enhances initial NK-cell cytotoxicity and cytokine secretion, whereas chronic exposure leads to ex vivo NK dysfunction. In vivo, CD48-overexpressing Vκ*MYC tumors progress more slowly and extend host survival, while NK-cell depletion accelerates disease. These findings support a context-dependent role for CD48, potentiating acute NK responses while coexisting with chronic NK exhaustion, and suggest strategies to modulate CD48 for therapeutic benefit.

## Introduction

Multiple myeloma (MM) is a clonal malignancy of plasma cells that profoundly reshapes the bone marrow (BM) microenvironment^1,2^. A hallmark of MM progression is the subversion of immune surveillance, particularly by natural killer (NK) cells, which are critical effectors in recognizing and eliminating malignant plasma cells^3,4^. In healthy individuals, NK-cell activity is governed by a balance of activating and inhibitory signals, mediated through surface receptors that engage ligands on target cells; disruption of this balance in MM contributes to impaired cytotoxicity and cytokine secretion, facilitating tumor immune escape^5,6^.

CD48 is a glycophosphatidylinositol (GPI)-anchored glycoprotein broadly expressed on hematopoietic cells that serves as a key ligand for CD244 (2B4), an NK-cell receptor containing immunoreceptor tyrosine-based switch motifs (ITSMs)^7^. Our prior surface proteomic analyses identified CD48 as one of the most abundantly expressed plasma membrane proteins on MM cell lines and primary MM tumors^8^. Moreover, CD48 has been described as a potential immunotherapeutic target in myeloma^9^, though its expression is not restricted to plasma cells^10^. Recent “structural surfaceomics” studies, combining crosslinking mass spectrometry (XL-MS) with surface biotinylation to identify tumor-specific protein conformations^11^, suggest that CD48 may adopt cancer-specific three-dimensional conformations in MM, potentially influencing its interactions with NK cells^12^.

Engagement of CD244 by CD48 can transmit either activating or inhibitory signals to NK cells, depending on downstream adaptors such as SLAM-associated protein (SAP), Ewing sarcoma–associated transcript 2 (EAT-2), and inhibitory phosphatases including SHP-1/2 and SHIP-1^7,13^. It is known that acute CD48–CD244 interactions can potentiate NK-cell activation, and in vitro functional genomics screens by us and others suggest CD48 to be a major NK-cell activator in MM^14,15^. However, emerging evidence suggests that persistent CD48 engagement may contribute to NK-cell dysfunction and exhaustion within the tumor microenvironment through dysregulation of SAP/SHIP-1^10,13,16^. Furthermore, as we show below, increased tumor *CD48* expression is associated with poor clinical outcomes in myeloma patients. Therefore, these observations frame a paradox: does high CD48 expression on MM plasma cells promote tumor elimination by NK cells, hindering tumor growth, or does it contribute NK cell dysfunction and/or exhaustion, allowing for tumor proliferation?

Despite the critical role of CD48 in MM immunosurveillance, the specific transcriptional circuits that drive *CD48* expression in malignant plasma cells, and how these circuits relate to NK-cell signaling, have yet to be fully elucidated. Furthermore, the causal links between CD48 abundance on MM cells, NK-cell receptor engagement, and downstream exhaustion signatures remain to be defined. To address these gaps, we combined bulk RNAseq patient-derived datasets, genome-wide CRISPR-based genetic screens, scRNA-seq profiling, in vitro NK–tumor co-culture assays, and in vivo experiments using an immunocompetent syngeneic Vκ*MYC myeloma model engineered for luminescence-based tumor tracking and CRISPR mediated gene editing. We implicate Estrogen Related Receptor Alpha (ESRRA) and Transcription Factor AP-2 Alpha (TFAP2A) as transcriptional regulators of *CD48* in MM cells, define NK-cell exhaustion signatures in RNAseq data from patient samples, and evaluate how modulating CD48 on tumor cells influences NK-cell activation in vitro, NK-cell function ex vivo, and tumor progression in vivo, findings validated by scRNA-seq analysis of NK cells from the Vκ*MYC model. Together, these results suggest that strategies to modulate CD48 could enhance NK-cell–mediated control of MM, particularly when combined with approaches to prevent or reverse exhaustion such as checkpoint inhibition.

## RESULTS

### CD48 Expression and Unsupervised NK-Ligand Clustering Stratify MM Patients by Cytogenetic Risk and Survival

To determine the landscape of *CD48* expression in MM and its clinical relevance, we analyzed bulk RNA-seq data from CD138+ malignant plasma cells in the Multiple Myeloma Research Foundation (MMRF) CoMMpass (IA19) cohort^17^. In this dataset, *CD48* expression was elevated in several high-risk cytogenetic subgroups. Specifically, t(4;14) and gain 1q21 cases exhibited significant (p < 0.0001) *CD48* upregulation relative to others, whereas t(11;14) (standard-risk) patients showed decreased *CD48* (p < 0.0001). When stratified by *CD48* expression, high *CD48* levels in tumors were associated with significantly lower overall survival (*p* = 0.0044) (**Fig. 1A**). These initial findings suggest the hypothesis that high MM tumor CD48 expression may be associated with reduced NK-cell immunosurveillance, enabling tumor growth.

**Figure 1.**
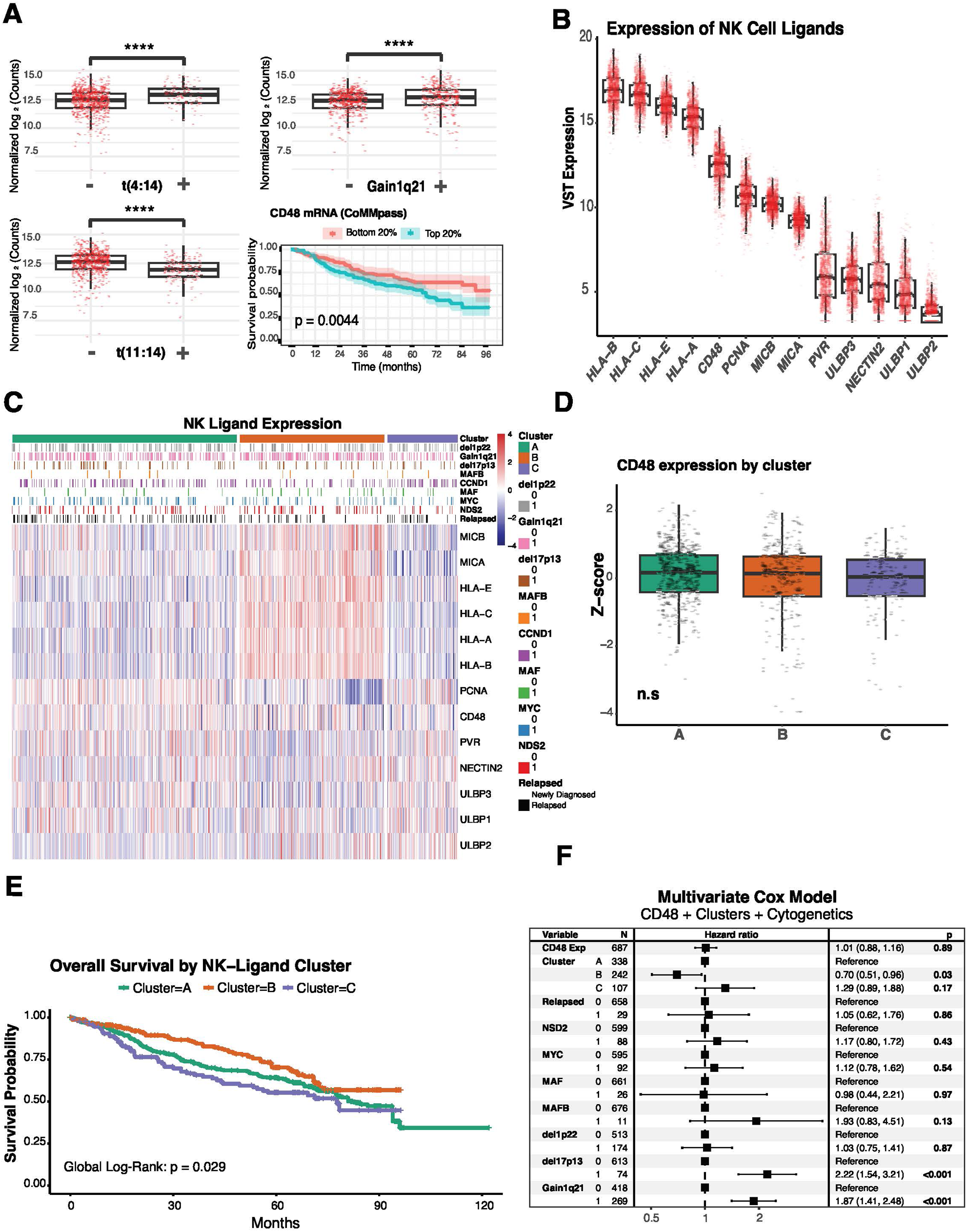
*CD48* Expression and Unsupervised NK-Ligand Clustering Stratify MM Patients by Cytogenetic Risk and Survival. **A.** Boxplots display normalized log₂ *CD48* mRNA expression in CoMMpass (IA19) CD138⁺ samples stratified by cytogenetic subgroup. Kaplan–Meier overall survival curves stratified by *CD48* expression (top 20% vs. bottom 20%) in 774 patients; log-rank. **B.** Boxplots of VST-normalized expression for a panel of NK-cell ligands across 774 MM samples. **C.** Heatmap of scaled expression for the same NK-ligands (rows) in all 774 CoMMpass patients (columns). Patients are arranged into three clusters: A (teal), B (orange), C (purple). Sidebars annotate key cytogenetic features: t(4;14), t(11;14), MAF, del(17p13), MAFB, NSD2, relapse status, MYC gain, gain(1q21). **D.** Boxplots comparing *CD48* log₂ expression among Clusters. **E.** Kaplan–Meier curves for overall survival by NK-ligand cluster. Global log-rank across A, B, and C: p = 0.029. Pairwise log-rank p-values B vs C = 0.026; B vs A = 0.085; A vs C = 0.221. **F.** Forest plot from a multivariate Cox proportional hazards model. Variables on the left include *CD48* expression (high vs. low), cluster membership, relapse status, and cytogenetic risk factors. Hazard ratios (black squares) with 95% confidence intervals are plotted on a logarithmic scale.

Given the hypothesis that high CD48 expression may contribute to NK cell dysfunction in MM, we next asked whether *CD48* upregulation occurs in the broader context of altered NK-cell ligand expression on tumor cells. To address this, we applied Variance Stabilizing Transformation (VST) to normalize expression values and quantified a panel of established NK cell ligands, *HLA-A/B/C/E*, *MICA/B*, *NECTIN2, PCNA*, *PVR*, *ULBP1–3* ^18^, in addition to *CD48*, across the same MM cohort. As expected, classical HLA class I genes (*HLA-A, -B, -C*) ranked highest in mRNA expression; notably, *CD48* was the next most abundant ligand on MM cells (**Fig. 1B**). These mRNA-based findings are consistent with high surface protein abundance of these antigens as found in our previous proteomic studies^8^. Hierarchical clustering of scaled ligand expression values revealed three distinct patient clusters: Cluster A exhibited intermediate expression across all NK ligand markers, along with high expression of PCNA. Cluster B was characterized by high expression of HLA-A, HLA-B, HLA-C, HLA-E, and MICA/B, while Cluster C showed low HLA class I expression and elevated levels of CD226 ligands (NECTIN2, PVR) and ULBP2 **(Fig. 1C; Fig. S1A)**. Notably, *CD48* expression was comparable across all three clusters (**Fig. 1D**). We further found these NK-relevant clusters correlated with key cytogenetic and clinical features. Cluster A was significantly enriched for CCND1 translocations (p < 0.001), gain 1q21 (p < 0.01), NSD2 status (p < 0.05), and relapsed disease (p < 0.01). In contrast, Cluster B showed an over-representation of MAF-translocated patients (p < 0.05) **(Table 1).** Kaplan-Meier analysis showed that overall survival differed across NK- ligand clusters (global log-rank across A, B, and C, p = 0.029). In pairwise tests, Cluster B outperformed Cluster C (B vs C, p = 0.026), with a nonsignificant trend versus Cluster A (B vs A, p = 0.085); Cluster A and Cluster C did not differ (p = 0.221).(**Fig. 1E**). In a multivariate Cox regression, including cluster membership, *CD48* expression, relapsed status, and key cytogenetic variables, assignment to Cluster B remained independently protective (p = 0.03), whereas *CD48* expression alone did not retain significance (p = 0.89). As anticipated, del(17p13) and gain(1q21) (both p < 0.001) were independently associated with poorer survival **(Fig. 1F).**

Collectively, these data establish that CD48 is elevated in high-risk MM subgroups and that NK-ligand expression patterns (particularly HLA class I) define patient clusters with distinct cytogenetic features and prognoses. In particular, the features defining Cluster B are associated with superior survival, underscoring that a broader NK-ligand landscape may be associated with anti-myeloma immunity and influences clinical outcomes. Furthermore, the results of our multivariate analysis suggest that *CD48* expression alone is not an independent driver of poor prognosis, setting the stage for further investigation of its function in MM.

### Combined Chromatin Profiling, Machine Learning, and Functional Screens Identify Candidate Regulators of CD48 Expression in Myeloma

Motivated to understand what regulates *CD48* expression in MM tumors, we first analyzed previously published ATAC-seq and ENCODE ChIP-seq profiling data^19^ from purified MM plasma cells, compared with normal donor plasmablasts and plasma cells. This analysis identified two MM-enriched accessibility peaks spanning the *CD48* promoter region (±1 kb of the transcription start site). Transcription factor (TF) footprinting analysis of ATAC-seq combined with ChIP-seq experimental data implicated 89 TFs as potential *CD48* regulators in primary MM tumors (**Fig. 2A**). Leveraging the CoMMpass dataset, we then trained an eXtreme Gradient Boosting (XGBoost) regression model to predict CD48 expression based on transcription factor (TF) activity. The model explained approximately 40% of the variance in CD48 expression (R² = 0.40; **Fig. 2B**). Feature importance was interpreted using SHAP (SHapley Additive exPlanation) values, which quantify each TF’s contribution to the predicted CD48 level across samples (positive SHAP values indicate genes whose higher expression predicts higher CD48 expression, whereas negative values indicate the opposite, **Fig. 2C**). Notably, IRF4, a well-known driver of myelomagenesis ^20–22^, was among the strongest positive predictors of CD48, followed by TFAP2A, previously identified by our group as a regulator of CD70 in MM ^23^, another prognostic surface protein. Conversely, MEF2B emerged as a potential negative regulator. Correlation analyses in CoMMpass confirmed these associations: CD48 and IRF4 showed a moderate positive correlation (R = 0.42, p < 0.0001), whereas CD48 and MEF2B showed a weak negative correlation (R = –0.096, p = 0.0035) (**Fig. 2D–E**).

**Figure 2.**
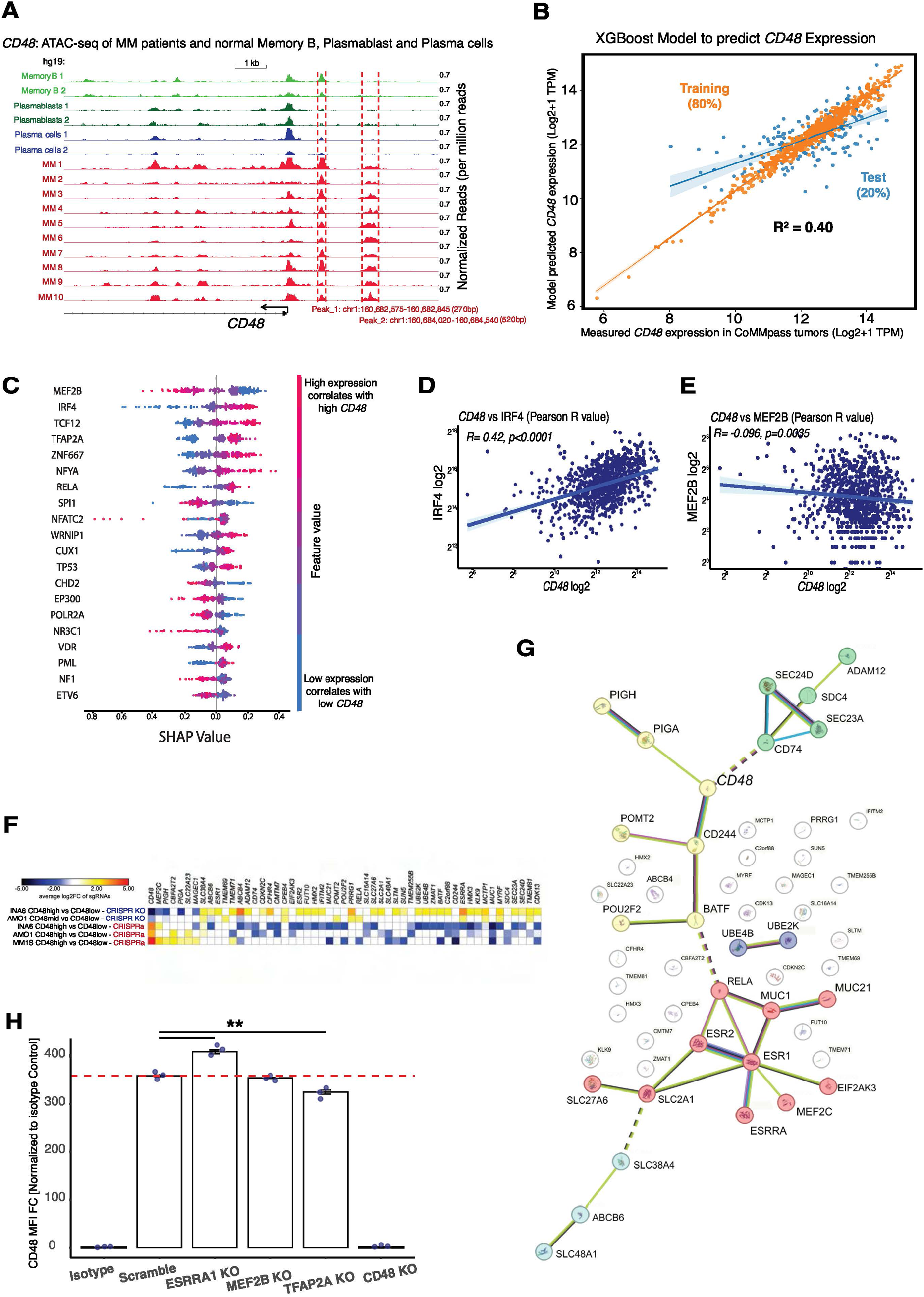
Combined Chromatin Profiling, Machine Learning, and Functional Screens Reveal Novel CD48 Regulators in Myeloma. **A.** ATAC-seq tracks at the *CD48* locus. Signal tracks display normalized read density (per million reads) from ATAC-seq of purified normal memory B (green), plasmablasts (blue), and plasma cells (purple), overlaid with ATAC-seq from primary MM samples (red). Two regions of enriched chromatin accessibility spanning ∼±1 kb around the *CD48* transcription start site (TSS) are highlighted with dashed red boxes. **B.** XGBoost predictive model for *CD48* expression. Scatter plot of observed *CD48* log₂ TPM (x-axis) versus model-predicted *CD48* (y-axis) in CoMMpass samples. Orange points and regression line represent the 80% training set; blue points and line represent the 20% test set. Model performance is indicated by R² = 0.40. **C.** SHAP (SHapley Additive exPlanation) summary plot identifying top transcription factors. Each dot represents a single patient’s SHAP value for a given TF (y-axis) plotted against the feature’s impact on *CD48* prediction (x-axis). TFs are ordered by mean absolute SHAP value; dots in pink (right) denote high SHAP values (positive impact on *CD48*), whereas blue dots (left) indicate negative impact. **D-E.** Correlation of *CD48* versus *IRF4* and *MEF2B* expression in CoMMpass. Scatter plot of CD48 log₂ versus *IRF4* log₂ TPM (y-axis). **F.** Heatmap showing average log₂ fold-change (log₂FC) of sgRNAs targeting the indicated genes in CRISPR KO (blue labels) and CRISPRa (red labels) genome-wide screens. Comparisons were performed between CD48 high and CD48 low populations across INA6, AMO1, and MM1S myeloma cell lines. Yellow indicates sgRNAs enriched in each condition, whereas blue indicates sgRNAs depleted. Data represent the top-ranked candidate genes consistently identified across replicates. **G.** STRING network of top surface CD48-modulating genes. Protein–protein interaction network generated by STRING for the 52 overlapping CRISPR-KO/a hits. Nodes are colored by functional cluster membership: nuclear steroid receptor activity (Red), GPI-anchor biosynthesis (Yellow), ER–Golgi transport (green), heme transport (light-blue), and ubiquitin ligase activity (purple). **H.** Flow-cytometric validation of CD48 surface expression following individual CRISPR-Cas9 knockout of *ESRRA*, *TFAP2A*, *MEF2B*, and *CD48* in AMO1 cells. Bars represent mean fluorescence intensity (MFI) fold change normalized to isotype control (n = 3 biological replicates; mean ± SEM). Asterisks denote statistical significance. Lack of asterisks indicates no significant difference.

To complement the machine learning predictions focused on transcriptional regulators of *CD48*, we determined additional regulators of surface CD48 through genome-wide CRISPR knock-out (CRISPR-KO) and activation (CRISPRa) FACS-based screens. These functional genomic screens across three MM cell lines (AMO1, INA6, and MM.1S) yielded 52 genes that reproducibly modulate CD48 surface expression (**Fig. 2F**). STRING network analysis of these hits organized them into five clusters: (a) nuclear steroid receptor activity, (b) glycosylphosphatidylinositol (GPI)-anchor biosynthesis, (c) ER–Golgi transport vesicle membrane components, (d) heme transmembrane transport activity, and (e) ubiquitin–ubiquitin ligase activity (**Fig. 2G**). These groupings suggested that *CD48* regulation occurs at multiple levels, from transcriptional control to post-translational processing and membrane trafficking. Functional enrichment analysis using g:Profiler^24,25^ highlighted significant biological themes consistent with these clusters (**Fig. S1B**) suggesting that CD48 regulation involves transcriptional, post-translational, and membrane trafficking processes.

Based on the combined evidence from STRING network analysis, functional enrichment, CRISPR-KO/a screen-concordance and machine learning prediction, we selected for validation three top candidate regulators of surface CD48: Estrogen Related Receptor Alpha (*ESRRA*), Myocyte Enhancer Factor 2B/C (*MEF2C*/*MEF2B*), and Transcription Factor AP-2 Alpha (*TFAP2A*). These targets were subsequently validated through CRISPR-mediated knockout experiments in the AMO1 myeloma cell line performed in pooled populations rather than single-cell clones to minimize clonal variation and maintain a representative population context. For *ESRRA*, we achieved ∼58% knockout efficiency, confirmed at the genomic level using synthego ICE tool^26^ (**Fig. S1C**), which resulted in a significant increase in surface CD48 expression compared to AMO1 scramble sgRNA (p < 0.01) (**Fig. 2H**). These data support CRISPR-KO/a screens findings that *ESRRA* represses CD48 and align with STRING and enrichment analyses. For *TFAP2A*, we achieved approximately 75% knockout efficiency, which correlated with a modest but statistically significant reduction in surface CD48 expression measured by MFI (p < 0.01) (**Fig. 2H; Fig. S1D**), further validating its role as a positive regulator predicted by machine learning. In contrast, *MEF2B* knockout was inefficient (∼12%), resulting in only minimal decreases in surface CD48 levels (**Fig. 2H**). An sgRNA targeting *CD48* served as a positive control and demonstrated effective knockout (**Fig. S1E**).

Together, this data suggests ESRRA functions as a putative repressor of CD48, whereas TFAP2A promotes its expression; MEF2B also might modulate CD48 via distinct transcriptional and post-translational mechanisms.

### CD48 Modulation Influences NK-Cell Activation In Vitro

Having identified transcriptional regulators of CD48 expression, we next sought to determine the functional consequences of modulating CD48 levels on NK-cell activation using a controlled in vitro co-culture system. We generated Vκ*MYC murine myeloma lines^27,28^ with stable *Cd48* knockout (CD48KO) or *Cd48* overexpression (CD48OE) (**Fig. S2A**). We then employed KIL C.2, a Jagged2-overexpressing murine NK-cell line on a C57BL/6 background, capable of degranulation and cytokine secretion upon IL-2 stimulation and tumor co-culture^29^, to serve as effector (**Fig. 3A**). KIL C.2 cells were co-cultured with Vκ*MYC WT, CD48KO, and CD48OE targets at effector-to-target (E:T) ratios ranging from 30:1 to 1:10 for 24 hours. KIL C.2 cells exhibited baseline cytotoxicity against Vκ*MYC WT. Although CD48OE cells showed a trend toward higher lysis and CD48KO cells showed slightly reduced lysis compared with WT, none of these differences reached statistical significance (**Fig. 3B**). Intracellular cytokine staining of KIL C.2 cells (gated as CD3⁻/NK1.1⁺/NKp46⁺) after 24 hours of co-culture confirmed NK-cell activation across all conditions. Co-culture with CD48OE Vκ*MYC cells resulted in a higher IFN-γ production compared with CD48KO targets and a modest increase in TNF-α, consistent with enhanced activation at both effector-to-target (E:T) ratio of 30:1 or 10:1. In contrast, CD48KO targets induced minimal IFN-γ and TNF-α expression **(Fig. 3C)**.

**Figure 3.**
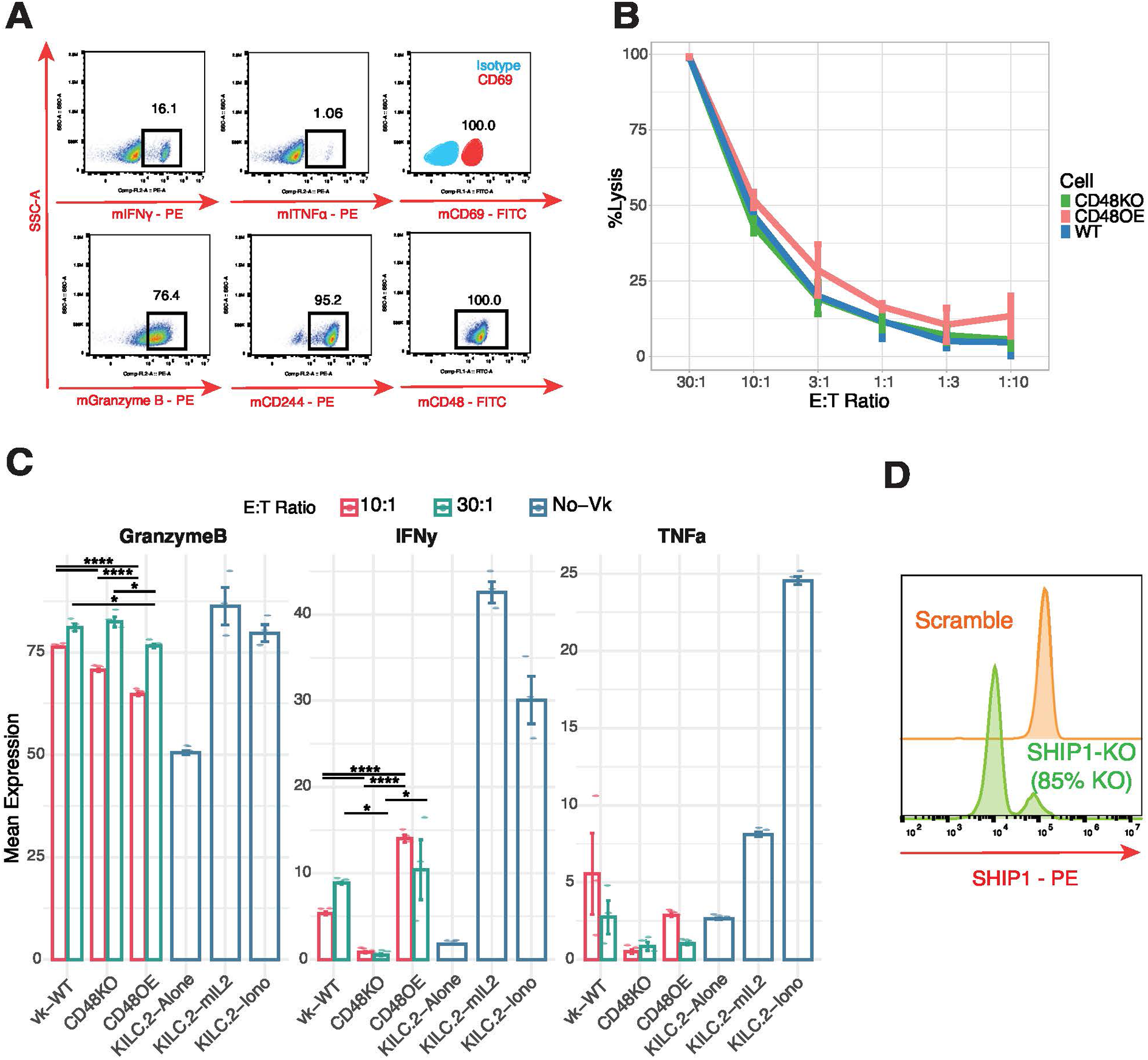
In Vitro NK–Tumor Co-Culture Functional Assays Demonstrate CD48-Dependent Activation and Limited Rescue by SHIP1. **A.** Representative flow-cytometry intracellular staining for murine IFN-γ, TNF-α, surface CD69, intracellular Granzyme B, surface CD244, and surface CD48 after co-culture with Vκ*MYC for 24 hours (E:T 30:1) (representative of n = 3 biological replicates). **B.** Percentage target-cell lysis of KIL C.2 against Vκ*MYC WT, CD48KO, and CD48OE across effector:target ratios. Points = mean ± SEM (n = 3 technical replicates). Pairwise Welch’s t-tests at each E:T (p-values; reported as WT–KO / WT–OE / KO–OE): 30:1 (0.494 / 0.548 / 0.954); 10:1 (0.295 / 0.184 / 0.077); 3:1 (0.902 / 0.422 / 0.413); 1:1 (0.967 / 0.510 / 0.183); 1:3 (0.582 / 0.436 / 0.613); 1:10 (0.876 / 0.351 / 0.373). No pairwise comparison reached p < 0.05. **C.** Bar graphs of mean KIL C.2 expressing Granzyme B, INFy and TNFa, measured at E:T = 30:1 (green circles), E:T = 10:1 (red circles), or “no-Vκ” PMA/ionomycin control (blue squares) (*n*=3 technical replicates). Statistical significance is indicated by asterisks; absence of asterisks denotes no significant difference (p ≥ 0.05). **D.** Flow cytometry histogram showing SHIP1 expression in KIL C.2 after CRISPR-Cas9 knockout of *Inpp5d* gene (representative of *n*=2 technical replicates).

To investigate whether SHIP1 contributes to NK-cell dysfunction in myeloma, we generated SHIP1KO KIL C.2 lines (*Inpp5d* gene knockout) using CRISPR-Cas9, confirmed by intracellular staining and Synthego ICE analysis (indel = 94%, R² = 0.95, Knockout-Score: 94) (**Fig. 3D; Fig. S2B**). SHIP1KO NK cells did not exhibit enhanced cytotoxicity against Vκ*MYC WT, CD48KO, or CD48OE targets at E:T ratios up to 30:1 after 24 hours (**Fig. S2C**), indicating that SHIP1 loss alone is insufficient to restore NK-cell function in this setting.

Together, these results indicate that higher CD48 expression on myeloma cells is associated with enhanced NK-cell activation, evidenced by increased cytotoxicity, cytokine production, and CD69 upregulation, whereas SHIP1 deletion in NK cells does not restore function. These findings indicate that CD48 expression influences NK-cell responses in myeloma, warranting further investigation of its immunoregulatory function to establish whether targeting CD48 could help reinvigorate NK-cell–mediated immunity in multiple myeloma.

### scRNA-Seq Reveals NK-Cell Exhaustion Signatures and Dysregulated CD48/CD244 Axis

To further explore *CD48* and *CD244* dynamics in myeloma primary tumors, we analyzed single-cell RNA-seq data from 8 healthy donors (HD) and 54 MM patients (GSE223060)^30^ (**Fig. 4A-B)**. To first examine the CD48-CD244 signaling axis, we compared expression across immune subsets from healthy donors (HD) and MM patients (**Fig. 4C**). A dot plot of *CD48* and *CD244* expression revealed that *CD48* is widely expressed across hematopoietic cells, in particular in macrophages and T-regs, with higher frequency and intensity in MM compared to HD (**Fig. 4D**). In contrast, *CD244* was mainly confined to NK cells and CD8+ T cells, with no major difference between MM and HD. To further interrogate CD48–CD244 interactions in the myeloma microenvironment, we applied CellChat^31,32^ analysis to infer ligand–receptor signaling between immune and malignant populations using this single-cell transcriptomes dataset selecting samples with 25% or more plasma cells. Circular and chord diagrams highlighted a strong directional flow of signaling from plasma cells, monocytes, macrophages, and B-cells toward NK cells **(Fig. 4F-G)**. Among all CD48-mediated interactions identified, the CD48–CD244 pair was significantly represented in the communication network, linking CD48⁺ myeloma plasma cells to CD244⁺ NK cells **(Fig. 4E)**. Although the overall interaction probability was lower than those between CD4⁺ T cells or macrophages and NK cells, the CD48–CD244 connection remained one of the most statistically significant ligand–receptor pairs within the CD48-associated network and not detected in HD samples by this method **(Fig. S3A)**, consistent with selective engagement in the myeloma microenvironment. Inferred ligand–receptor analysis further revealed that CD48–CD244 signaling ranked among the significant CD48-mediated interactions in MM, along with HLA-E and CD74 axis **(Fig. 4E**, **Fig. 4H),** which has also been described as key immune checkpoints in NK cells in myeloma^33^. Although the CD48–CD244 connection appears less prominent than some other predicted interactions, CellChat identified it among the statistically significant and most representative ligand–receptor pairs linking plasma cells to NK cells in the MM microenvironment. Importantly, this interaction was not detected in healthy donor samples, supporting its disease-specific relevance. Additional strong signaling routes were inferred from T cells and monocytes to NK cells, consistent with the broader immune activation context in MM. Overall, these findings indicate that CD48–CD244 communication is preferentially represented in myeloma and may contribute to altered NK-cell signaling within the tumor microenvironment.

**Figure 4.**
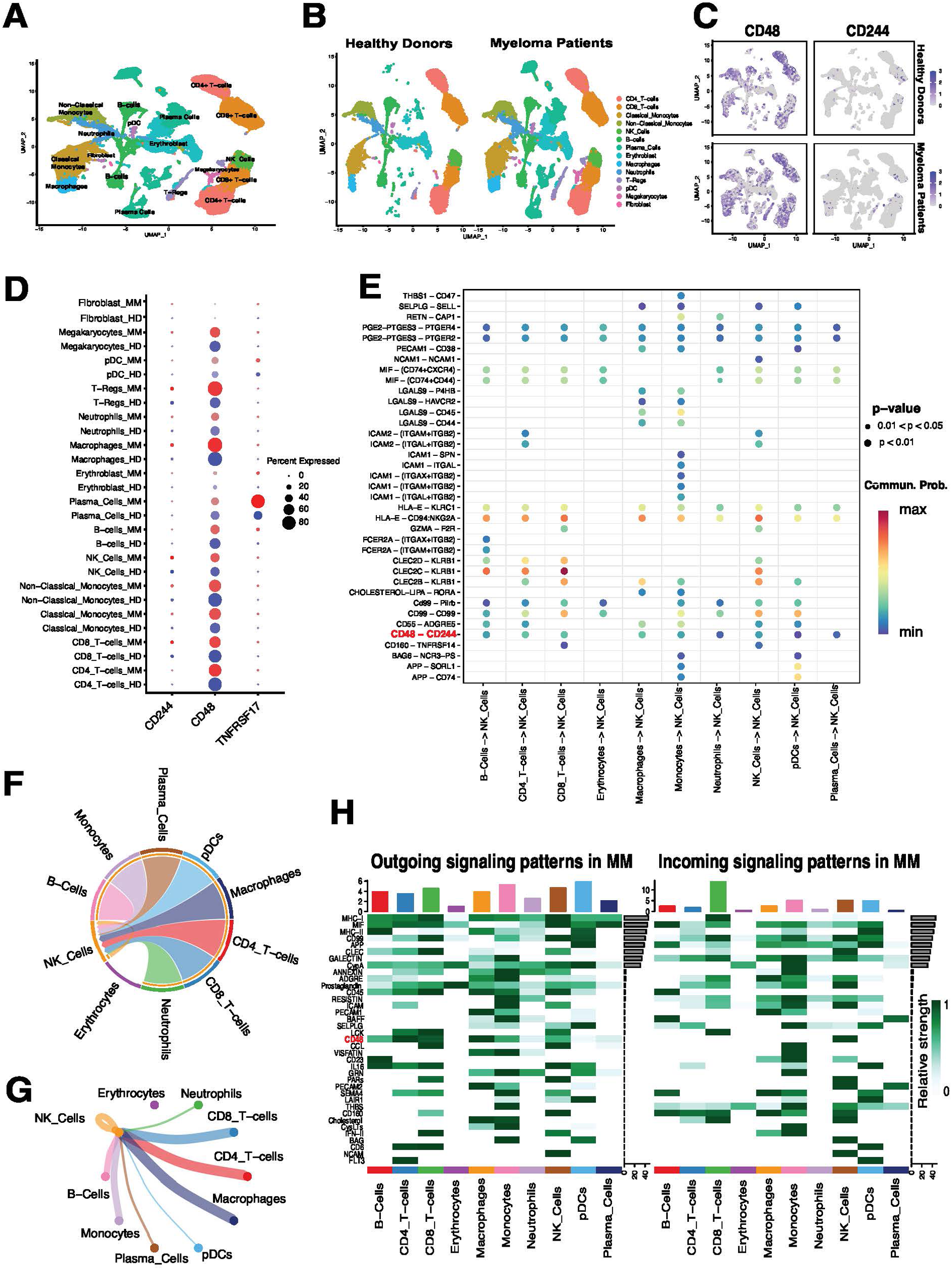
Single-cell RNA-seq and CellChat analysis reveal CD48–CD244 signaling between malignant plasma cells and NK cells in myeloma. **A.** UMAP visualization of cell types identified across healthy donors (HD) and multiple myeloma (MM) samples based on scRNA-seq data (GSE223060). **B.** UMAP split by condition (HD vs. MM), showing shared immune populations across disease states. **C.** Feature plots showing expression of *CD48* and *CD244* across all cells. **D.** Dot plot showing expression levels and percent-expressing cells for *CD48*, *CD244*, and the plasma cell marker *TNFRSF17*, split by condition (Blue HD and red MM). **E.** Dot plot of individual ligand–receptor pairs predicted by CellChat to interact with NK cells. Circle size represents statistical significance (permutation test p-value) and circle color indicates communication probability. Among these, CD48–CD244 is highlighted as a significant and strong predicted interaction, suggesting it is one of the dominant NK-cell communication axes in MM. **F.** Chord diagram generated using CellChat, illustrating CD48 signaling pathway network from plasma cells, monocytes, macrophages, and B-cells, predominantly targeting NK-cells. **G.** Circle plot showing CD48 signaling pathway network depicting directional CD48 signaling interactions, highlighting strong engagement from CD48⁺ immune and malignant cells to CD244⁺ NK cells. **H.** Heatmaps of outgoing and incoming signaling strength across cell types in MM, as inferred by CellChat. Interaction strength reflects the relative contribution of each signaling pathway to intercellular communication. The CD48–CD244 axis is highlighted within this broader network of significant signaling pathways.

Building on this predicted cell–cell interaction network, we next examined how sustained CD48–CD244 engagement might contribute to the altered transcriptional and functional state of NK cells in MM. Compared to HD-NK cells, MM-NK cells exhibited significantly higher expression of the inhibitory phosphatase SHIP1 (*INPP5D*; Wilcoxon, p = 0.0012) and the activating phosphatase SAP (*SH2D1A*; Wilcoxon, p = 0.0029) (**Fig. 5C**). These findings suggest that phosphatase-mediated signaling is altered in the myeloma microenvironment, indicative of NK cell re-education. Within the NK compartment, we identified MM-enriched subclusters displaying an “CMV adaptive-like” CD56–dim phenotype^34,35^ (**Fig. 5A-B**; **Fig. S3F**), corresponding to the recently described NK3 subset^36^. This subcluster was characterized by high expression of cytotoxic effector genes (*GNLY*, *PRF1*, *NKG7*, *FGFBP2*) (**Fig. 5B**) and receptor repertoire shifts: significant downregulation of *KLRB1* and *KLRF1*, alongside upregulation of *KLRC2* and *KLRC3*. (**Fig. S3B**).

**Figure 5.**
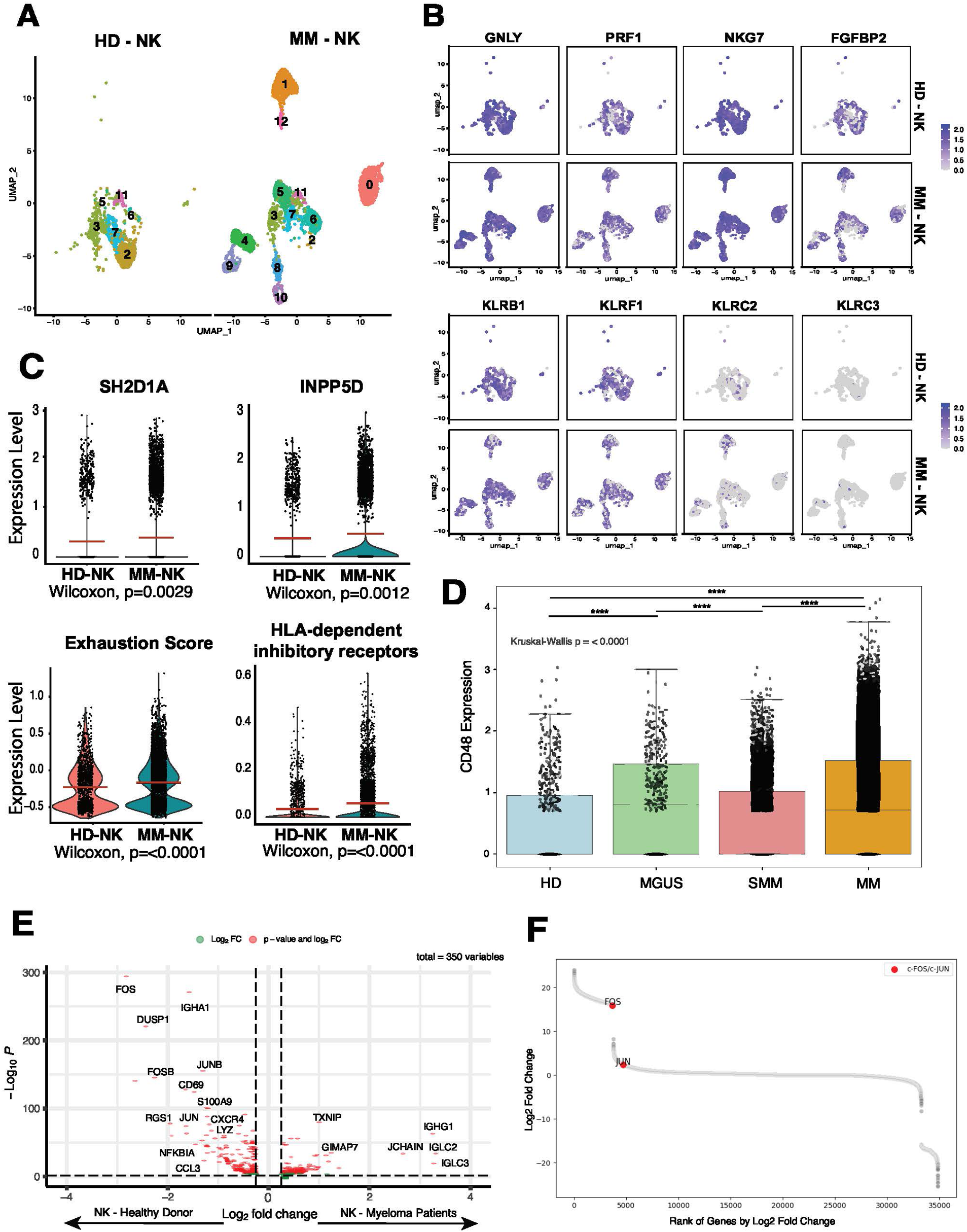
Single-Cell RNA-Seq Reveals NK-Cell Exhaustion Signatures and Dysregulated CD48-CD244 Axis in Human Multiple Myeloma. **A.** UMAP of NK-cell subclusters only from 8 healthy donors (HD) and 54 MM patients from GSE223060 (HD-NK on left; MM-NK on right). **B.** UMAP feature plots of key NK-cell cytotoxicity and receptor genes on the top row. Bottom row: receptor genes related to adaptive CMV CD56dim phenotype. **C.** Violin plots compare expression and calculated scores in HD-NK (orange) versus MM-NK (green). **D.** Boxplot of CD48 expression in plasma cells across four groups: Control (Blue), MGUS (green), SMM (Pink), MM (Orange) from >200 samples from 182 individuals multiple myeloma, MGUS, and SMM, and non-cancer controls^42^. Global Kruskal–Wallis: *p* < 0.0001. Dunn’s post-hoc pairwise tests: HD–MGUS: p= 3.37×10^-13^; HD–SMM: p= 0.193; HD–MM: p= 2.82×10^-26^; MGUS–SMM: p= 3.34×10^- 15^; MGUS–MM: p= 0.540; SMM–MM: p=1.90×10^-141^. Asterisks above boxes denote significance (**** *p*<0.0001; *** *p*<0.001; ** *p*<0.01; * *p*<0.05; ns, not significant). **E.** Volcano plot of differential gene expression in NK cells (MM vs. HD) from GSE223060. **F.** Rank plot of gene expression fold changes in HD-NK versus MM-NK (HD > 0, MM <0) from a bigger MM and HD cohort (MM n=61, HD n =91) ^42^.

Further underscoring NK cell dysfunction in MM, an exhaustion module score, calculated via SCENIC AUCell^37^ using a published gene set^35,38^, was markedly higher in MM-NK than in HD-NK (Wilcoxon *p* < 0.0001) (**Fig. 5C**). In parallel, HLA-dependent inhibitory receptors were significantly upregulated in MM-NK (Wilcoxon *p* < 0.0001), whereas HLA-independent inhibitory checkpoints showed no significant difference (*p* = 0.74) (**Fig. 5C**, **Fig. S3D**).

Differential expression analysis (HD vs. MM) further revealed that AP-1 family members, c-Fos (*FOS*), c-Jun (*JUN*), JunB (*JUNB*), FosB (*FOSB*), and *CD69* were markedly downregulated in MM-NK (log₂FC range: –1.5 to –2.8; adjusted p < 0.0001) (**Fig. 5E**). In contrast, *BATF* was relatively enriched in MM-NK (Wilcoxon p < 0.0001) (**Fig. S3C**), consistent with its known role as a core transcription factor and key driver of NK cell dysfunction in other hematological malignancies such as acute myeloid leukemia (AML)^39^. Loss of AP-1 activity has been linked to impaired NK maturation, reduced cytokine production (IFN-γ, TNF-α), and diminished NKG2D expression, mirroring phenotypes seen in murine myeloid leukemia and exhausted CAR-T models^40,41^.

To validate these AP-1 findings across larger cohorts, we examined a publicly available scRNA-seq dataset containing >200 samples from 182 individuals with MM, MGUS, SMM, and non-cancer controls^42^. Here, c-Jun (*JUN*) and c-Fos (*FOS*) were upregulated in non-cancer control NK cells versus MM-NK, confirming AP-1 downregulation in MM (**Fig. 5F**). Additionally, *CD48* expression was highest in MM samples compared to the precursor conditions of SMM or MGUS, and healthy donors (**Fig. 5D**).

Overall, these data demonstrate that MM-NK cells display a combination of heightened activating receptor expression and co-occurring exhaustion signatures, characterized by increased inhibitory receptor and phosphatase expression, AP-1 downregulation, and reduced CD69, reflecting a dysfunctional state in the myeloma microenvironment likely driven by chronic stimulation. Although this environment includes CD48–CD244 engagement among many MM-specific ligand–receptor interactions not observed in healthy donors, the transcriptional features described here do not establish CD48 as a unique or direct mechanistic driver of these NK-cell changes.

### NK-Cell Depletion Accelerates MM Progression in Vκ*MYC Mouse Model

To assess tumor engraftment and the immunocompetent microenvironment, we employed the syngeneic Vκ*MYC model (Vκ32245 line)^28^, a non-secretory myeloma line we lentivirally transduced with Firefly luciferase (effLuc) and mCherry to enable in vivo tracking. To evaluate potential immunogenic rejection, we compared engraftment of Vκ*MYC effLuc/mCherry in NSG versus BL6 mice (**Fig. S4A**). We additionally engineered a Vκ*MYC effLuc/mCherry line to express dCas9-KRAB-mTagBFP2 for CRISPRi, together with a doxycycline-inducible sgRNA vector^43^. As proof of concept, CRISPRi activity was validated by targeting murine CD138 (mCD138) and confirmed by flow cytometry (**Fig. S4E**). In BL6 mice, effLuc signal emerged approximately one week later than in NSG controls, both in the EffLuc-only and EffLuc-dCas9 lines. This delay may reflect a combination of immune surveillance effects and reduced bioluminescence sensitivity due to darker skin and hair pigmentation in the C57BL/6 background, while still demonstrating successful engraftment and in vivo tracking **(Fig. S4A–B)**.

After confirming that Vκ*MYC cells engraft and emit luminescence in BL6 mice, we injected 1 × 10⁶ effLuc-tagged Vκ*MYC WT cells intravenously into C57BL/6 mice. NK cells were depleted with anti–NK1.1 (clone PK136; 200 µg i.p. twice weekly) or an isotype control (**Fig. 6A**). Flow cytometry of peripheral blood (CD3neg/NK1.1pos/NKp46pos) confirmed >98% NK-cell depletion by Day 7, which persisted through Day 42 (**Fig. 6B, Fig. S4C**). Bioluminescence imaging showed accelerated tumor growth in NK-depleted mice by Day 21 compared to isotype-treated controls (**Fig. 6C and 6D**). Correspondingly, NK-depleted mice had shorter median survival **(Fig. S4D)**.

**Figure 6.**
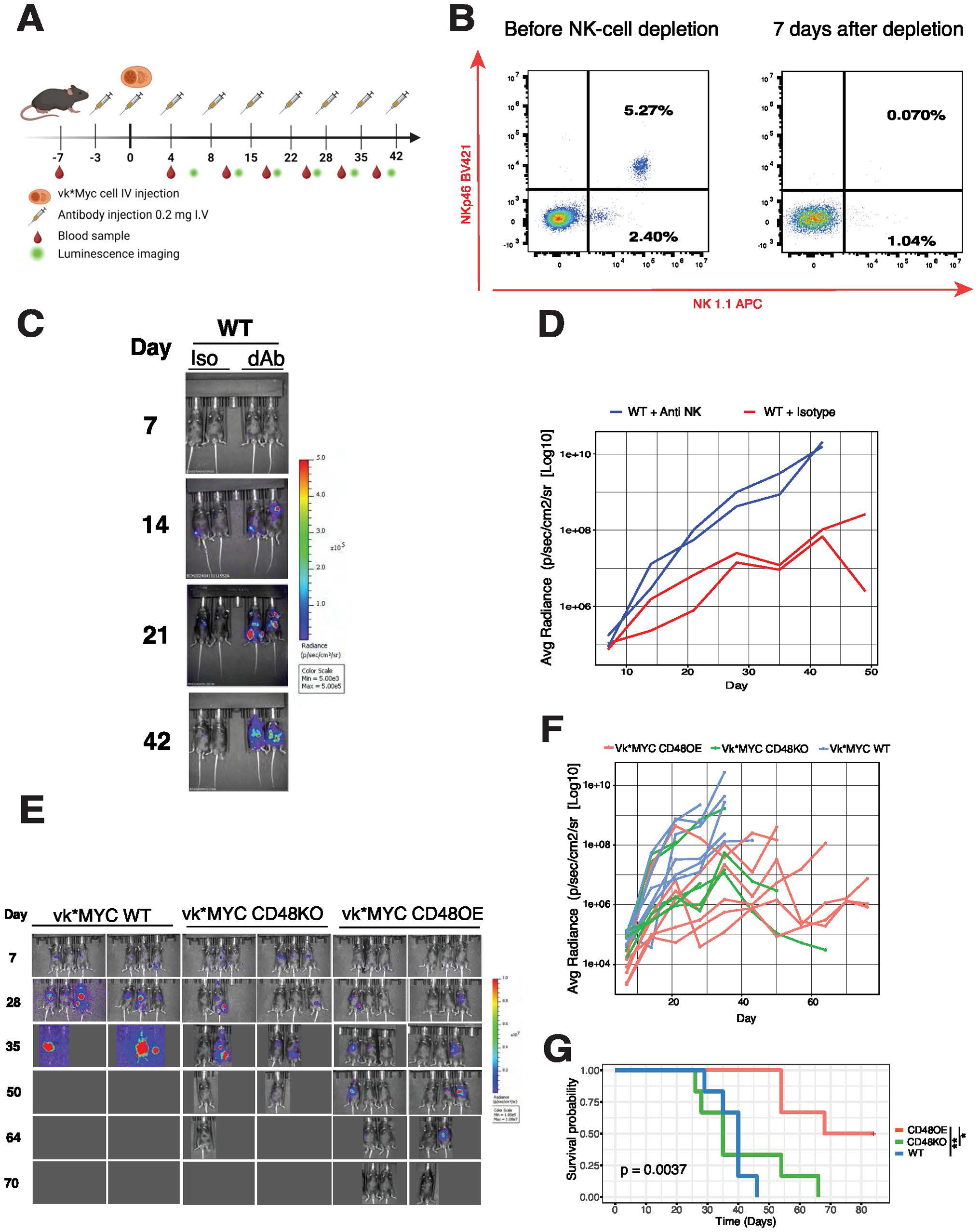
In Vivo Vκ*MYC NK-Cell Depletion and CD48 Overexpression Impact Tumor Engraftment and Survival. **A.** Experimental timeline and schematic of NK-cell depletion and IgG control using Vκ*MYC WT injected into C57BL/6 mice. **B.** Representative flow-cytometry plots for NK-cell detection on peripheral blood. NK cells (NK1.1+/NKp46+) before and after depletion. **C.** Serial BLI images of Vκ*MYC WT tumors in BL6 mice under NK depletion antibody (dAb) or Isotype control (Iso), (n=2 per arm). Color scale indicates radiance (photons/sec/cm²/sr). **D.** Quantification of BLI signal over time for Vκ*MYC WT + anti NK1.1 antibody or IgG Isotype control. **E.** Representative BLI images for WT, CD48KO, and CD48OE (n=6 per arm). **F.** Individual longitudinal BLI trajectories for each mouse bearing Vκ*MYC WT, CD48KO, or CD48OE cells (n=6 per arm). **G.** Kaplan–Meier survival curves for mice injected with Vκ*MYC WT, CD48KO, or CD48OE cells tested by log-rank test.

These results confirm that the Vκ*MYC model is a robust system with reliable tumor tracking via Firefly luciferase–based bioluminescence, while the CRISPRi-engineered line offers a versatile tool for functional studies. Importantly, NK cell depletion in this immunocompetent model led to accelerated disease progression, highlighting the critical role of NK cells in controlling myeloma.

### CD48 overexpression in Vκ*MYC myeloma cells slows disease progression and prolongs survival

Given the critical role of NK cells in controlling myeloma progression, we next investigated whether modulating tumor-intrinsic CD48 expression could impact engraftment dynamics and disease outcome in vivo.

When 1 × 10⁶ CD48OE cells were injected intravenously into naïve C57BL/6 mice (n = 6 per arm), engraftment occurred by Day 14, but subsequent tumor progression was significantly slower than in WT or CD48KO, despite similar initial engraftment (**Fig. 6E-F**). Correspondingly, median survival extended from 40 days (WT) to 68 days (CD48OE; log-rank p = 0.0037) (**Fig. 6G**). CD48KO tumors exhibited engraftment kinetics similar to WT, likely reflecting the low endogenous CD48 expression in WT (**Fig. S2A**).

Together, these results support that CD48 overexpression on Vκ*MYC cells slows tumor progression and prolongs survival in immunocompetent mice. Although CD48OE line engrafted at the same time as controls, it exhibited markedly slower growth, translating into extended median survival. These findings confirm that CD48 overexpression impairs tumor progression in vivo, underscoring CD48’s role in modulating anti-myeloma immunity.

### Ex Vivo NK-Cell Functional Impairments Mirror Transcriptomic Exhaustion

As CD48 overexpression altered tumor progression, we next examined whether it also influenced NK cell functional status. Splenic and bone marrow NK cells were isolated at terminal disease from Vκ*MYC tumor–bearing mice (WT, CD48KO, CD48OE) and naïve controls and assessed ex vivo by plate-bound stimulation (**Fig. 7A-B**).

**Figure 7.**
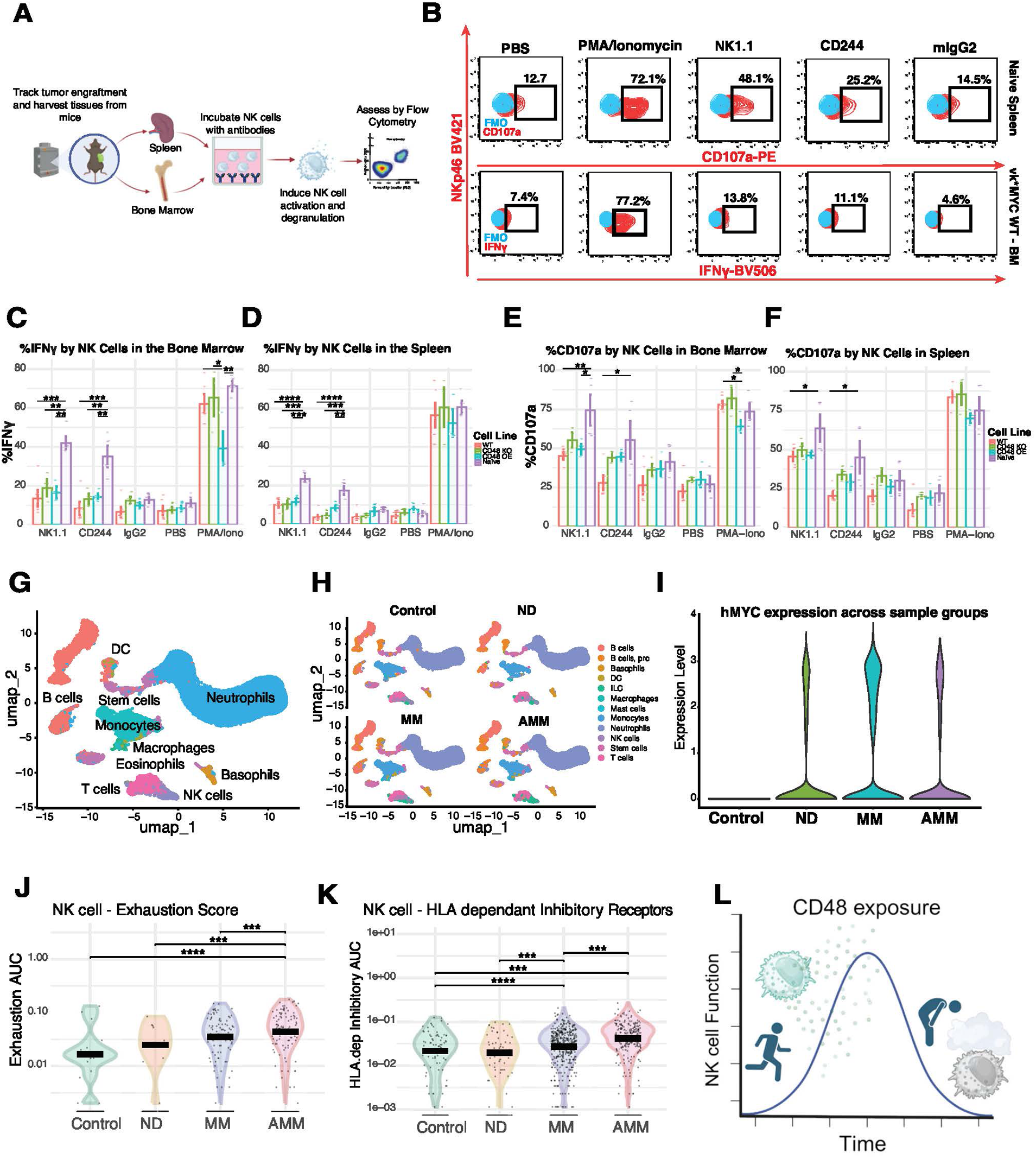
Ex Vivo NK-Cell Functional Impairments and scRNA-Seq Dynamics in Vκ*MYC Model. **A.** Workflow schematic for ex vivo NK-cell functional assays. Spleens and bone marrow (BM) are harvested from Vκ*MYC-bearing (WT, CD48KO, CD48OE) and naïve control mice. Single-cell suspensions of NK-cells are incubated in high-binding 96-well plates coated with specific antibodies (anti–NK1.1, anti–CD244), isotype control (mIgG2a), or PBS negative controls for 6 hours. After stimulation, NK-cell activation and degranulation are assessed by flow cytometry (intracellular IFN-γ, surface CD107a). **B.** Representative flow-cytometry gating for ex vivo NK activation in Spleen and Bone Marrow. **C-F.** Splenic and Bone Marrow NK-cell IFN-γ secretion and CD107a degranulation. Bar graphs show mean IFN-γ median fluorescence intensity (MFI) and CD107a MFI in spleen and bone marrow-derived NK cells from naïve controls, Vκ*MYC WT, CD48KO, and CD48OE groups after stimulation with anti–NK1.1, anti–CD244, isotype (mIgG2a), PBS, or PMA/ionomycin (naïve n = 4; tumor-bearing groups n = 6 per genotype). **G.** UMAP projection of scRNA-seq data from Vκ*MYC BM samples (GSE134370). Major hematopoietic cell populations are color-coded and labeled. **H.** Split UMAPs showing 4 different subgroups and batch reduction. Control, newly diagnosed (ND), multiple myeloma (MM), and advanced MM (AMM) samples from the immunocompetent Vκ*MYC model. **I.** Violin plots of human MYC expression across four groups. **J-K.** Violin plots of exhaustion and HLA-Dependent inhibitory AUCell score for individual NK cells in four groups: Control, ND, MM, AMM. **L.** CD48 exposure initially promotes NK cell activation and degranulation, but high CD48 cannot prevent longer-term exhaustion.

NK cells from tumor-bearing mice exhibited marked impairment in IFN-γ secretion and CD107a upregulation in response to anti–NK1.1 or anti–CD244 stimulation, regardless of CD48 status, compared to robust responses in naïve NK cells (**Fig. 7C–F**). These defects were consistent across both spleen and bone marrow, indicating a broad functional impairment consistent with NK cell dysfunction (**Fig. S5A-B**).

Notably, even under PMA/ionomycin stimulation, CD48OE-derived NK cells trended toward lower cytokine output than WT or CD48KO groups, suggesting a potential additive exhaustion phenotype in the myeloma microenvironment in the syngeneic Vκ*MYC model (**Fig. 7C–F**).

Together, these data demonstrate that NK cells from tumor-bearing mice are functionally exhausted ex vivo, regardless of CD48 status on Vκ*MYC, highlighting the profound immunosuppressive effects of the myeloma niche.

### scRNA-Seq Data Reveal NK-Cell Functional Dynamics Across Myeloma Progression in the Vκ*MYC Model

Finally, to complement our experimental studies above, we leveraged scRNA-seq data from a *de novo* Vκ*MYC mouse cohort across stages of disease progression (GSE134370)^44^ in the C57BL/6/KaLwRij background (Control: n = 3, early-MM (ND): n = 5, intermediate-MM (MM): n = 3; advanced-MM (AMM): n = 7) (**Fig. 7G-I**), with stages defined by serum M-protein levels. We calculated eight NK-cell module scores (cytotoxicity, inflammatory response, stress response, HLA-dependent inhibitory receptors, HLA-independent inhibitory receptors, HLA-dependent activating receptors, HLA-independent activating receptors, and exhaustion).

Although cytotoxicity module scores were overall low across all groups, NK cells from MM and AMM mice showed a modest but statistically significant reduction compared with controls **(Fig. S5D)**, whereas ND mice showed no difference relative to controls. In contrast, the HLA-independent activating receptor score increased progressively from Control to MM, and AMM **(Fig. S5C)**, suggesting an inflammatory NK-cell phenotype in the absence of effective tumor control. Exhaustion scores were significantly elevated in MM compared to Control and rose further in AMM **(Fig. 7J)**. Finally, HLA-dependent inhibitory receptor expression was elevated in both MM and AMM groups relative to controls (**Fig. 7K**).

Overall, NK cells show progressive functional impairment, characterized by loss of cytotoxicity, and rising exhaustion. Notably, these murine NK-cell transcriptomic profiles parallel key patterns observed in human MM scRNA-seq datasets, where NK cells in newly diagnosed and relapsed patients likewise exhibit stepwise increases in inflammatory and exhaustion signatures.

## DISCUSSION

Our study provides a comprehensive, multi-platform analysis of CD48’s role in regulating NK-cell immunity in multiple myeloma (MM)^14,15^. Integrating prior proteomic data with transcriptomic, genetic, and functional analyses, we find that CD48 is highly expressed in high-risk MM subsets and prominently positioned within the NK-ligand landscape associated with patient outcomes. CD48 emerged as the second-most highly expressed NK-ligand (after HLA class I) in the CoMMpass dataset; however, unsupervised clustering also highlighted composite NK-ligand expression patterns, particularly *HLA-A*/*B*/*C*/*E* combined with *MICA*/*B*, as relevant prognostic markers, and *CD48* alone was not an independent prognostic factor. This reveals a paradox: although CD48 functions as a critical NK-cell activator, its elevated expression correlates with poorer clinical outcomes. One possible contributing factor is the location of the CD48 gene on Chromosome 1q, which is frequently gained in high-risk MM and associated with high-risk features^45^. While our functional data suggest that CD48 upregulation enhances NK-cell recognition and surveillance, this immune visibility may be outweighed by the aggressive biology of CD48-high tumors. Alternatively, increased expression of activating NK cell ligands has been shown in other settings to promote hyporesponsiveness through chronic stimulation^46,47^. By analogy, sustained CD48 exposure in the MM microenvironment may initially stimulate increased NK cell surveillance of MM tumors but appears insufficient to prevent immune escape, and arises in parallel with NK-cell dysfunction and disease progression over time **(Fig. 7L)**. Our data are consistent with this model but do not establish CD48 itself as the driver of the exhaustion phenotype.

At the transcriptional level, our analyses pinpointed *ESRRA*, *TFAP2A*, and *MEF2B* as potential modulators of *CD48*, highlighting multiple control points from transcriptional initiation to post-translational trafficking. CRISPR-Cas9 validation experiments confirmed *ESRRA* and *TFAP2A* as regulators of CD48, whereas *MEF2B* knockout did not produce a detectable change compared with the scramble control. This likely reflects the low editing efficiency observed for *MEF2B* (∼12%) and baseline ploidy variation that can limit complete gene disruption in myeloma cells. Moreover, these transcription factors operate within partially overlapping regulatory networks, such that the loss of a single factor may be buffered by other co-expressed regulators. Together, these findings underscore the robustness of *CD48* transcriptional control in plasma cells. Functional enrichment consistently linked these candidates to steroid-receptor signaling, ER-associated processing, and GPI-anchor biosynthesis, suggesting that modulating CD48 in MM may require targeting not only transcription factors but also ER and GPI-anchoring machinery. Although ESRRA’s role in MM is not well established, high ESRRA activity in other cancers suppresses immune pathways, such as CD8⁺ T-cell recruitment and NK infiltration, potentially shielding tumor cells from immune detection^48,49^. This multi-checkpoint control of CD48 could, in principle, be exploited not only to fine-tune surface density but also to regulate its temporal expression, enabling more precise control of NK-cell engagement and potentially avoiding exhaustion.

scRNA-seq profiling of human MM samples revealed that NK cells adopt an “adaptive-like” CD56–dim phenotype with elevated SHIP1 and SAP phosphatases, downregulated AP-1 family members, and increased inhibitory receptors, collectively defining an exhausted transcriptomic signature. Although these NK cells upregulated both HLA-dependent and HLA-independent activating receptors, they remained dysfunctional, while downregulated AP-1 family mirrors observations in CML^40^, exhausted CAR-T cells^41^ and more recently other scRNAseq analysis^50^. The Vκ*MYC mouse model recapitulated many of these features: as disease progressed from early to intermediate and active MM, NK cells lost cytotoxicity, acquired inflammatory and stress signatures, upregulated HLA-dependent inhibitory/activating receptors, and displayed rising exhaustion. These parallel murine and human NK phenotypes validate the Vκ*MYC model as an informative platform for dissecting NK-cell dysfunction in MM.

Functional assays corroborated that CD48 on MM cells acutely enhances NK activation, but ex vivo analyses of NKs from tumor-bearing mice showed loss of degranulation and cytokine secretion upon receptor cross-linking. Even PMA/ionomycin stimulation failed to fully rescue cytokine production in CD48OE-exposed NKs. These results highlight a potential duality: while CD48 can potentiate NK effector responses initially, persistent CD48 expression in the MM microenvironment may be associated with NK-cell dysfunction.

In vivo, CD48 overexpression on Vκ*MYC cells slowed tumor progression and extended survival in immunocompetent mice. Conversely, NK-cell depletion markedly accelerated MM progression, underscoring NK cells’ protective role even when partially exhausted and illustrating how profound dysfunction accelerates disease and relapse. Together, these findings suggest that increased CD48 expression on MM cells may enhance early NK-mediated tumor control, whereas persistent expression coincides with features of an immunosuppressive microenvironment.

Both therapeutic and biological implications emerge from these findings. First, therapeutic strategies to transiently boost CD48, potentially via epigenetic or transcriptional modulators of ESRRA/TFAP2A, could sensitize MM to NK-cell clearance, particularly if paired with interventions that prevent or reverse exhaustion (for example, NK checkpoint blockade or metabolic reprogramming). Second, our data emphasize that MM treatment approaches must consider the dynamic NK-ligand landscape: focusing solely on CD48 may be insufficient if concurrent HLA or MICA/B expression dampens NK-cell activity. Notably, Cluster B features (characterized by elevated expression of *HLA-A*/*B*/*C*/*E* and *MICA*/*MICB*), suggest that robust CD8⁺ T-cell activity and NKG2D-driven NK-cell activation may counterbalance inhibitory signaling from KIR/HLA and NKG2A/HLA-E. This interplay between activating and inhibitory pathways likely contributes to the improved survival observed in Cluster B, highlighting the importance of the overall NK-ligand environment beyond CD48 alone in shaping anti-myeloma immunity and influencing clinical outcomes. Rational combination therapies, increasing CD48 while preserving or restoring AP-1 function and blocking HLA-E/NKG2A interactions could synergize to restore anti-myeloma immunity. Our data underscore the need for therapeutic strategies that carefully modulate CD48 to restore effective NK-cell surveillance without exacerbating exhaustion.

While our study integrates multiple platforms to define the immunoregulatory role of CD48 in MM, several limitations should be acknowledged. First, although the Vκ*MYC model and the KIL C.2 NK-cell line capture essential aspects of MM biology, they may not fully represent the complexity of human disease, or the diversity observed in primary NK cells. Nevertheless, our scRNA-seq analysis of NK cells from the Vκ*MYC model across distinct disease stages closely mirrors findings from human MM patient samples. This similarity underscores the translational relevance of our findings and demonstrates that the Vκ*MYC model also effectively recapitulates critical aspects of immune dysfunction observed in human MM. Notably, to our knowledge, we have also generated the first stable CRISPRi-expressing Vκ*MYC model, which provides a valuable new tool for functional studies and could serve as a resource for the broader myeloma research community.

Additionally, our approach to CD48 overexpression utilized an immature, non-cleavable GPI-anchored isoform, limiting exploration into the potential roles of soluble CD48 isoforms. Soluble forms have been described for other NK cell immune ligands, including CD226 ligands (NECTIN2, PVR) and ULBP2 (predominantly found in Cluster C, which is associated with the worst prognosis in our analysis), where they are thought to modulate immune interactions^18^. Future validation studies using patient-derived xenograft or humanized models will therefore be essential to fully capture human immune complexity and rigorously evaluate therapeutic strategies targeting CD48. Importantly, our Vκ*MYC results support the notion that CD48 primarily functions to engage NK cells and enhance anti-tumor immunity in vivo. However, determining whether CD48 itself (versus other MM-associated ligands) plays a central role in the NK-cell exhaustion phenotype we observe will require additional mechanistic work. Furthermore, several of our transcriptomic and ligand–receptor interaction findings, particularly those based on CellChat and scRNA-seq, would benefit from additional validation using primary NK cells from MM patients and complementary functional assays.

Overall, our study provides novel insights into the role of CD48 in MM. By integrating proteomic, genomic, and functional approaches, we show that elevated CD48 expression enhances NK-cell activation, particularly in early disease contexts. In advanced MM, NK cells display pronounced exhaustion features, but our data do not support a direct role for CD48 in preventing or driving this dysfunctional state. Instead, CD48 appears as one component of a broader, MM-specific NK-ligand network, that differs from the HD setting, within which we identify key transcriptional and post-translational regulators (including ESRRA, TFAP2A, and MEF2B) as promising therapeutic targets. Importantly, our findings suggest that strategies designed to transiently increase CD48 availability or modulate its upstream regulators may enhance NK-cell–mediated antitumor responses without worsening exhaustion. Together, this work offers mechanistic insights and provides a rationale for developing combination immunotherapies aimed at restoring NK-cell function and improving outcomes for patients with multiple myeloma.

## Contributions

B.P.E. and A.P.W. designed and conceived the study. B.P.E. and T.S. performed experiments and analyzed data. B.P.E., L.P.L., and S.R. performed experiments. O.A.A., and C.S.M. participated in experimental design. B.P.E. and H.G. analyzed patient datasets. B.P.E and D.W. analyzed murine datasets. F.S., P.P., J.A.C.S., Y.Z., and V.S. performed murine studies. B.P.E. and A.P.W. wrote the manuscript. All authors provided feedback and approved the manuscript.

## METHODS

### Cell lines

Human myeloma cell lines (MM.1S, AMO-1, INA-6) were obtained from ATCC or DSMZ, authenticated by STR analysis, and routinely tested for mycoplasma. They were cultured in RPMI 1640 medium supplemented with 20% FBS, GlutaMAX^TM^, and 100 U/mL penicillin-streptomycin. The murine Vκ*MYC myeloma cell line (Vκ32245) was kindly provided by Drs. Marta Chesi (Mayo Clinic, Scottdale, AZ) and maintained in the same medium. The murine NK cell line KILC.2 (Kerafast®) was cultured in IMDM supplemented with 30% FBS, GlutaMAX, penicillin-streptomycin, and β-mercaptoethanol, with added murine SCF (50 ng/mL, Peprotech, 250-03) and IL-7 (25 ng/mL, Peprotech, 217-17). For cytotoxicity assays, KILC.2 cells were pre-activated with murine IL-2 (20 ng/mL, Peprotech, 212-12) for 24 hours prior to co-culture.

### Molecular Cloning and DNA Plasmids

Gene fragments encoding murine CD48 (mature and immature isoforms; UniProt P18181), and dCas9-KRAB with a BFP2-mTag reporter were synthesized by Twist Bioscience (South San Francisco, CA). Constructs were assembled into lentiviral expression vectors using Gibson assembly (NEB, E2611S) and verified by plasmid sequencing (Plasmidsaurus, South San Francisco, CA). Plasmids were amplified in *E. coli* NEB 5-alpha Competent Cells (High Efficiency; NEB, C2987H) and Stbl3 Competent Cells (Macro Lab, UC Berkeley, CA). DNA was purified using either the QIAprep Spin Miniprep Kit (Qiagen, 27104) or the Plasmid Plus Midi Kit (Qiagen).

### Lentiviral Vector Production

Lenti-X 293T cells (Takara, Cat. #632180) were transfected with plasmids encoding murine CD48, firefly luciferase–mCherry, or dCas9–BFP2-mTag using the TransIT™-Lenti Transfection Reagent (Mirus Bio, Cat. #MIR6600), according to the manufacturer’s instructions. Cells were cultured for 2–3 days post-transfection, and lentiviral supernatants were collected at 48 and 72 hours. Viral particles were concentrated using Lenti-X Concentrator (Takara Bio, Cat. #631232) following the manufacturer’s protocol.

### Flow Cytometry

Cells were resuspended in FACS buffer (D-PBS containing 2% FBS) and stained with fluorescently conjugated antibodies for 20–40 minutes at 4 °C. After staining, cells were washed twice and resuspended in fresh FACS buffer. Samples were acquired on a CytoFLEX (Beckman Coulter), or Cytoflex Flow Cytometer (Beckman Coulter, Beckman Coulter Navios Flow Cytometer) or FACSAria-Fusion or FACSAria III flow cytometer (BD Biosciences). Data were analyzed using FlowJo software (version 10.10.0). Fluorescence compensation was performed using UltraComp eBeads™ (Invitrogen, Cat. #01-2222-42). For murine samples analysis, gating strategy used was FSC-A/SSC-A for lymphocyte population, single cells gated in SSC-A/SSC-H and then NK cell population was gated on murine CD3-/NK1.1+/NKp46 to determine IFN-γ, CD107a, CD69, CD48 and CD244 expression.

### In vitro cytotoxicity assays

The Vκ*MYC cell line (Vκ32245) was stably transduced with luciferase and engineered to express various CD48 densities. For cytotoxicity assays, 50,000 tumor cells were seeded in IMDM-cell media in white 96-well plates (Greiner Bio-One), followed by co-culture with pre-activated KILC.2 cells at varying effector-to-target (E:T) ratios (30:1 to 1:10) in IMDM, with triplicate wells per condition. After 18–24 hours of incubation at 37 °C, D-luciferin (375 μg/mL, Gold Biotechnology, LUCK-1G) was added, and luminescence was measured using a GloMax Explorer (Promega). Bioluminescence signals were averaged across replicates and normalized within each condition to generate a 0–100% cytotoxicity scale.

### Murine studies

All animal experiments were conducted in accordance with UCSF Institutional Animal Care and Use Committee (IACUC) guidelines. NSG (NOD.Cg-Prkdc^scid Il2rg^tm1Wjl/SzJ) and C57BL/6J mice (6–9 weeks old, both sexes; Jackson Laboratories) were bred and housed at the UCSF Preclinical Therapeutics Core. Our study examined male and female animals, and similar findings are reported for both sexes. Mice were injected intravenously via the tail vein with 1 × 10⁶ Vκ*MYC (Vκ32245) tumor cells stably expressing firefly luciferase. Tumor burden was monitored weekly using bioluminescence imaging (IVIS, PerkinElmer), beginning 7 days after tumor injection. Humane endpoints were determined based on signs of illness and in accordance with veterinary protocols. For NK cell depletion ^51^, mice received intravenous injections of anti-NK1.1 antibodies (Bio X Cell; clone PK136, Cat. #BE0036) or isotype control (mouse IgG2a, Cat. #BE0085) at 0.2 mg per dose on days –3, 0, 4, and 8, relative to tumor challenge. From day 8 onward, depletion was maintained with weekly dosing.

### Murine blood NK cell detection

Peripheral blood was collected via submandibular vein puncture using sterile lancets. Samples were obtained weekly during the first four weeks after initiation of the NK-cell depletion protocol and every other week thereafter. Peripheral blood was collected from all mice in each experimental group at each time point unless otherwise indicated. Red blood cells were lysed using ammonium chloride lysis buffer (ACK buffer), and the remaining leukocytes were washed and resuspended in D-PBS supplemented with 2% FBS prior to flow cytometric analysis. NK cells were identified using antibodies against murine CD3, NK1.1, and NKp46. All samples were analyzed on a CytoFLEX flow cytometer (Beckman Coulter) as described above.

### Plate-Bound Stimulation Assay for Murine NK Cells

To evaluate NK cell activation via receptor engagement^52^, 96-well high-binding plates (Corning Costar Brand 96-Well EIA/RIA Plates, Cat: 07200721) were coated overnight at 4°C or 2 hr at 37°C with PBS-diluted antibodies (typically 25 µg/mL; e.g., anti-NK1.1 [PK136]). Control wells contained PBS or PMA/ionomycin. After washing, single-cell suspensions from murine spleen and bone marrow were added without prior NK enrichment (0.5–1 × 10⁶ cells/well in complete media). Anti-CD107a and monensin were added, and cells were incubated for 6 hours at 37°C. Post-stimulation, cells were stained for viability, NK markers (NK1.1, NKp46, CD3ε), and intracellular IFN-γ. Flow cytometry was used to assess degranulation (CD107a) and cytokine production.

### CRISPRa/KO Genome-wide Screens

Multiple myeloma cell lines were engineered to stably express either SpCas9 (for knockout studies) or dCas9-VP64 (for activation studies). Cells were transduced at low multiplicity of infection with genome-scale sgRNA libraries (Brunello for CRISPR KO [Addgene #73179], Calabrese for CRISPRa [Addgene # 1000000111]) and cultured under puromycin selection to maintain library representation. Following recovery, cell pools were stained for surface CD48 (APC anti-human CD48 Antibody, Biolegend #33714) and sorted by fluorescence-activated cell sorting (FACS, FACSymphony, S6, BD Biosciences) into CD48-high, CD48-mid and CD48-low populations. Genomic DNA was isolated, and integrated sgRNA cassettes were PCR-amplified and subjected to next-generation sequencing (NGS) to quantify sgRNA abundance in each sorted population. Sequencing reads were mapped to the corresponding library reference and analyzed using MAGeCK. For each gene, enrichment or depletion of sgRNAs in the CD48 high versus CD48 low fractions was assessed. Genes were considered significant if they displayed enrichment or depletion of ≥3 independent sgRNAs and achieved a MAGeCK p-value <0.05. For visualization, the average log₂ fold-change (log₂FC) of normalized read counts per gene was calculated. Genes that did not meet the significance criteria or had |log₂FC| <0.25 were collapsed to zero. Hits were further prioritized if they scored concordantly in at least two independent screens or showed functional relationships with other significant regulators.

### CRISPR-Cas9 Gene Knockouts

Cas9 protein and either target-specific or scrambled sgRNA (Synthego, Redwood City, CA) were complexed at a 1:2.5 molar ratio and incubated at 37 °C for 10–15 minutes. A total of 1 × 10⁶ Vκ*MYC, AMO1, or KILC.2 cells were washed with PBS and resuspended in 20 μL of nucleofection solution containing the Cas9–sgRNA complex. KILC.2 cells were electroporated using the P3 Primary Cell 4D-Nucleofector™ X Kit S and program CM-137; Vκ*MYC and AMO1 cells were electroporated using the SF Cell Line 4D-Nucleofector™ X Kit S and program DS-137 (Lonza). Immediately post-nucleofection, cells were plated into 80 μL of pre-warmed IMDM or RPMI 1640 media, incubated at 37 °C for 15 minutes, and then transferred to complete recovery media. CD48-negative clones were identified by flow cytometry and sorted using a FACSAria Fusion or FACSAria III (BD Biosciences). Human and murine sgRNA sequences used were taken from Brunello and Brie Library respectively^53^: Human: TFAP2A: AUCCUCGCAGGGACUACAGG, ESRRA: AGACACCAGTGCATTCACTG, CD48: ATGTACAGTG CGCCACTCTG, MEF2B: CCTCTTCCAGTATGCCAGCA, CD74: AGCCGC GGAGCCCTGTACAC, TCGCGCTGGTCATCCATGAC. Murine: Cd48: GAACGAGUUGAAGAUAACCC, Inpp5d (SHIP1): CAUCCGAAGGUGUCCCCAUG (sgRNA sequence from Synthego tool). Scrambled sgRNA: non-targeting control specified by Synthego.

### Genomic DNA confirmation of CRISPR-Cas9 knockouts

Genomic DNA was extracted from 1 × 10⁶ Vκ*MYC or KILC.2 cells (fresh or frozen) using the Monarch® Genomic DNA Purification Kit (NEB, Cat. #T3010S). Primers were designed using the IDT PrimerQuest™ Tool (Integrated DNA Technologies, Coralville, IA) to generate ∼500 bp amplicons, with at least 200 bp flanking each side of the sgRNA target site. Primers were reconstituted in nuclease-free water to a final concentration of 10 μM. PCR amplification was performed using NEBNext® High-Fidelity 2X PCR Master Mix (NEB, Cat. #M0541S) with 0.5 μM of each primer and 1 ng to 1 μg of genomic DNA. PCR products were purified using the QIAquick PCR Purification Kit (QIAGEN, Cat. #28104) and submitted for Sanger sequencing (Quintara Biosciences, Fort Collins, CO). Resulting .ab1 files were analyzed using the ICE (Inference of CRISPR Edits) tool from Synthego (https://ice.synthego.com), and indel percentage, model fit (R²), and knockout score were used to evaluate editing efficiency. Primer sequences: m*INPP5D (SHIP1).* Set 1: Forward: GGATGTGAGGCGTGAGTG, Reverse: GAAAGCATCCTTCAACCTCCA. Set 2: Forward: ATACTTCTATTTGCACCTTTGC, Reverse: CCTCCAAAGAGGTACTCTAGTC. Set 3: Forward: TTTGCAAAGGGCTCCTG, Reverse: TAGTCTAGAAGCACCCGTGA. m*CD48.* Set 1: Forward: GGACCCACTTGGACCATATAAA, Reverse: TAGGTCAGGCTCACTCTCAA. Set 2: Forward: GGCAGCAATGTAACCCTGAA, Reverse: GCAGCAGCTATCAGCTTTGA. Set 3: Forward: CCACCGGCAGCAATGTA, Reverse: GCTTTGATATCTGGTAGGTCAGG. MEF2B: Set 1: Forward: CATGTTCGAGGAAGGGTTAGG, Reverse: AGCCACAAAGCCAGATGAA; Set 2: Forward: TTACCAGCTCAGCCAGAAAG, Reverse: CAGCAGAAATAAAGCACGTCAG. CD244: Set 1: Forward: TCAGCTTGGGTTCTCTTTCC, Reverse: ATAGGACACATTGCCATCCC; Set 2: Forward: TGACCATGTGGTTAGCATCTC, Reverse: TGTACCAAGCATAGGACACATT. ESRRA: Set 1: Forward: CTCACAGCCTGTCCACAAA, Reverse: GACCACAATCTCTCGGTCAAA; Set 2: Forward: CTGGGAACAGGGACGATTT, Reverse: CAAAGAGGTCACAGAGGGTAG. TFAP2A: Set1: Forward: CCACCCTACCAGCCTATCT, Reverse: ACTCTCCGCCTCCGAATAA, Set 2: Forward: CCCAATGCCGACTTCCA, Reverse: CTGTATGTTCCAGGTATCCTTTCT.

### CRISPRi sgRNA Cloning and Lentiviral Transduction for a Doxycycline inducible Knock-down system in Vκ*MYC

To generate sgRNA-expressing constructs for CRISPR interference (CRISPRi), we used the FgH1tUTG lentiviral vector (Addgene #70183), which harbors an H1 promoter with a Tet-inducible sgRNA cassette^43^. Guide sequences (∼20 bp) were selected from the Weissman lab CRISPRi-v2 library^54^ and modified with 4-bp overhangs (5′-TCCC for the forward strand and 5′-AAAC for the reverse complement) to allow directional cloning into the BsmBI-v2-digested FgH1tUTG backbone. The vector was linearized by overnight digestion with BsmBI-v2 at 55°C, followed by gel purification. Forward and reverse oligos were annealed in T4 ligase buffer by heating to 95°C for 2 minutes and slowly cooled to room temperature. Annealed oligos were diluted (1:100 final) and ligated into the dephosphorylated backbone using T4 DNA ligase with a 1:20 vector-to-insert molar ratio, as calculated via the NEBioCalculator. Ligation reactions were incubated at 16°C overnight, and 1–5 µL of the ligation product was transformed into competent *E. coli*. Lentiviral particles were produced and used to transduce dCas9-KRAB-BFP2mTag Vκ*MYC (Cas9-KRAB–expressing). Transduced cells were sorted based on fluorescent reporters (e.g., BFP and GFP) for downstream phenotypic or functional assays. For treatment of Vκ*MYC cells to induce expression of the sgRNA, doxycycline hyclate(Sigma-Aldrich D9891) was dissolved in sterile water at a stock concentration of 50 mg/ml and added to tissue culture medium for a final concentration of 1 µg/ml for 48-72 hours, before phenotypic confirmation by flow cytometry. Oligos used to clone sgRNA sequences used for CD138 knock down were as follows: **<u>TCCC</u>**GGAGACAGAGCCTAACGCAG, **<u>AAAC</u>**CTGCGTTAGGCTCTGTCTCC. Scramble sgRNA: **<u>TCCC</u>**GGGAACCACATGGAATTCGA, **<u>AAAC</u>**TCGAATTCCATGTGGTTCCC

### CoMMpass cytogenetic subtype transcriptional analysis

RNA-seq data from primary multiple myeloma (MM) samples in the MMRF CoMMpass study (release IA19) were downloaded via the MMRF Researcher Gateway. Differential expression analysis based on cytogenetic risk and CD48 expression was conducted using DESeq2 (v1.34.0) ^55^ in R (v4.1.2), with LFC shrinkage applied via the *apeglm* method ^56^. Functional enrichment and gene namespace conversion were performed using *gprofiler2* (v0.2.2)^24^. Cytogenetic classifications were assigned using associated metadata to compare CD48-high and CD48-low groups. Survival analyses were performed with the *survminer* (v0.4.9) and *survival* (v3.5-7) packages, comparing the top and bottom 20% of CD48-expressing patients. Correlations between CD48 expression and predicted transcriptional regulators were visualized using *ggscatter* from the *ggpubr* package (v0.6.0).

### CoMMpass analysis for NK cell ligand clustering in tumor cells

Gene expression data from the MMRF CoMMpass study were analyzed to profile NK cell ligand expression in multiple myeloma. Raw counts were normalized using variance-stabilizing transformation (VST) from DESeq2, and Ensembl IDs were mapped to HGNC symbols. A curated panel of NK ligands (e.g., MICA, CD48, ULBPs, HLA genes) was extracted and visualized using boxplots and heatmaps. Unsupervised hierarchical clustering of ligand expression defined distinct tumor sample groups, which were annotated with cytogenetic alterations and clinical stage (newly diagnosed vs. relapsed). Cluster–feature associations were assessed via Fisher’s exact test, and survival outcomes were evaluated using Kaplan-Meier analysis.

### Multivariate Cox Proportional Hazards Modeling for CD48

Differential gene expression analysis based on cytogenetic risk was performed on RNAseq data from primary MM samples from the Multiple Myeloma Research Foundation (MMRF) CoMMpass study (release IA19). Sample-level cytogenetic metadata, including translocations and copy-number alterations (del17p13, gain1q21, and del1p22(NSD2, MYC, MAF, MAFB), along with NK cell cluster groups, were extracted from the provided SeqFISH files and merged by sample identifier. Samples missing any metadata or with non-intersecting identifiers were excluded. Using clinical and survival information in CoMMpass, we fit a Cox model using the survival package^57^ (v3.8-3) for cytogenetic variables, and relapse status was coded as factors (levels 0/1). Hazard ratios (HRs) and 95% confidence intervals were extracted from the model summary. Statistical significance was assessed at α = 0.05, two-sided. We visualized the multivariate HRs using the forestmodel package (v0.6.2)^58^, and composed with ggplot2 (v3.5.1) with all covariates displayed on a single forest plot^59^. The Cox model was fit by maximizing the partial likelihood (with Efron’s method for ties), and inference was provided via the Wald test for individual coefficients and the likelihood-ratio and score (log-rank) tests for the overall model fit.

### ChIP-seq and ATAC-seq analysis

Raw ATAC-seq and H3K27Ac ChIP-seq data from MM patients and normal Memory B cells, Plasmablasts, and Plasma cells were downloaded from the European Nucleotide Archive (accession: PRJEB25605). Paired-end reads were aligned to the human reference genome (hg19) using Bowtie2. Only uniquely mapped reads with ≤2 mismatches were retained. Clonal reads, defined as reads mapping to the same genomic position and strand, were collapsed into a single representative read. Peak calling was performed using MACS2 or HOMER program. To investigate transcriptional regulation of CD48, motif enrichment analyses were conducted within ATAC-seq peaks located near the CD48 promoter. Motif predictions were generated using the PROMO tool and ENCODE transcription factor databases (V3 builds 336 and 161).

### Machine learning analysis for transcriptional regulation of CD48 expression

A curated list of 89 transcription factors was generated from ATAC-seq, ChIP-seq, PROMO, and ENCODE databases for model construction. RNA-seq data (TPM values) from 776 CD138⁺ myeloma samples in the CoMMpass MMRF dataset (release IA19) were log-transformed and used as input features. An XGBoost (Extreme Gradient Boosting) model was implemented using the *xgboost* package in Python via Google Colab. All transcription factors co-expressed with CD48 were included as predictors. Hyperparameter optimization was performed using randomized search with 5-fold cross-validation across 5,000 iterations, exploring parameters including colsample_bytree, colsample_bylevel, colsample_bynode, gamma, learning_rate, max_depth, n_estimators, and subsample. Model interpretability was assessed using SHAP (Shapley Additive Explanations) analysis, applied to the test set (20% of data). Model interpretability was assessed using SHAP (SHapley Additive Explanations) analysis applied to the test set (20% of data). SHAP values quantify each transcription factor’s contribution to the model’s predicted CD48 expression for each individual sample, with positive SHAP values indicating that higher expression of the factor predicts higher CD48 expression, and negative values indicating the opposite. Mean absolute SHAP values were used to rank transcription factors by their overall importance, and summary plots (Fig. 2C) were generated to visualize these relationships, where each dot represents one patient and color indicates the relative expression of each factor (red = high, blue = low). Model performance was evaluated using R² and adjusted R² values, as well as mean absolute error across folds to ensure robustness across training/test splits. This approach parallels our previously published analyses of transcriptional regulators for CD38, CD70, and CD46 in MM ^23,60,61^.

### scRNA-seq Analysis of CD48, CD244, and NK Cell Function

Single-cell RNA sequencing data from GSE223060 ^30^ was analyzed using Seurat (v4.3.0) ^62^ in R (v4.1.2). Cells with fewer than 500 detected genes, fewer than 1,000 total RNA counts, more than 50,000 features, or >10% mitochondrial gene content were excluded. Normalization and identification of highly variable genes were performed prior to integration. Batch correction was conducted using the Harmony package (v0.1.1)^63^, grouping by experimental condition. Dimensionality reduction and clustering were performed using UMAP (resolution = 0.5). Gene symbols were annotated using biomaRt, and cluster annotation was informed by prior publication metadata ^30,35^. Visualization was performed with ggplot2^64^ (v3.5.1) and gridExtra (v2.3)^65^.

### Differential Gene Expression and AUCell Analysis in NK Cells

Within NK cells, differential gene expression (DGE) analysis comparing MM vs. control and other pairwise groups were conducted using Seurat’s FindMarkers() (Wilcoxon test) on log-normalized data. AUCell (v1.16.0)^37^ was used to calculate per-cell AUC enrichment scores for curated gene sets reflecting various NK cell functional states. Genes were ranked per cell and AUC scores were derived from the top-expressed genes. The same curated gene sets (human or murine orthologs as appropriate) were used in both human and murine analyses ^38^: Cytotoxicity: GZMA, GZMB, GZMH, GZMK, PRF1, CTSW. Inflammatory cytokines/chemokines: CCL3, CCL4, CCL5, CXCL8, CXCL10, IL2, IL6, IL7, IL15, IL18. Stress/immediate early response: FOS, FOSB, JUN, JUND, EGR1, HSPA1A, HSPA1B, HSP90AA1, SOCS3, DUSP1, IER2, ZFP36. HLA-dependent inhibitory receptors: LAG3, LILRB1, LILRB2, KIR2DL1, KIR2DL3, KIR3DL1, KIR3DL2. HLA-independent inhibitory receptors: PDCD1, TIGIT, HAVCR2, CD96, CD300A, SIGLEC7, SIGLEC9. HLA-dependent activating receptors: KIR2DS4, KIR3DS1, CD160. HLA-independent activating receptors: NCR1, NCR2, NCR3, KLRK1, CRTAM, FCGR3A.Exhaustion markers: CISH, LAG3, TIGIT, KLRG1, KLRC1, PDCD1. AUC scores were stored in the Seurat metadata and visualized with violin plots across disease groups. Wilcoxon rank-sum tests with FDR correction were used to assess statistical differences.

### Cell-Cell Communication Analysis with CellChat

To explore ligand–receptor signaling in MM bone marrow, CellChat (v1.6.1) ^32^ was applied to a subset of MM patients from the integrated Seurat object (patients with >25% plasma cells). Data were extracted from the “data” slot of the RNA assay. Cell type labels were based on Seurat identities. Using CellChatDB.human, overexpressed genes and interactions were identified and signaling networks were inferred with communication probabilities. Networks were filtered (minimum 10 cells per group), and CD48-specific interactions were visualized using circle and chord diagrams as well as heatmaps with netVisual_aggregate() and netVisual_heatmap().

### Validation Using Public scRNA-seq Compendium

A large public collection of scRNA-seq and scTCR-seq data from more than 200 samples from 182 patients with multiple myeloma, MGUS, and SMM, and non-cancer controls^42^ (panImmune.h5ad), was accessed through Zenodo (https://zenodo.org/records/13646014). Data was primarily collected from Gene Expression Omnibus (GEO) Maura et al. under accession GSE161195^66,67^, Bailur et al. (GSE163278 ^68^). Oetjen et al. (GSE120221^69^). Granja et al. (GSE139369^70^), Zavidij et al. (GSE124310^2^), Kfoury et al. (GSE143791^71^), and Zheng et al. (GSE156728^72^). Analysis was done in Python (Scanpy v1.9) via Google Colab on an A100 GPU. Sample group annotations (“HD,” “MGUS,” “SMM,” “MM”) were assigned based on regex matching of sample IDs. After QC filtering (<500 genes, <1,000 RNA counts, >10,000 features, >10% mitochondrial content), standard Scanpy normalization and log1p transformation were applied. CD48 expression was visualized using Seaborn boxplots across groups.

### Murine Vκ*MYC scRNA-seq Analysis

Raw counts from GSE134370 ^44^ (murine Vκ*MYC model) were processed in Seurat (v4.3.0). After QC (nFeature_RNA >200 and <2,500, <5% mitochondrial content), data were normalized and batch-corrected using Harmony. Dimensionality reduction was done using PCA and UMAP. Cell types were annotated using SingleR with the ImmGen reference, and non-immune cells were removed. NK cells were subset and tumor contamination excluded based on expression of the human MYC transgene. AUCell was applied using the same functional gene sets (converted to murine orthologs) as the human analysis. Group comparisons (Control, ND, MM, AMM) were conducted using violin plots and Wilcoxon rank-sum tests with FDR correction.

## Code Availability

All custom scripts and code used for bioinformatic and statistical analyses, including bulk and single-cell RNA-seq analysis, survival modeling, machine learning, and visualization are available in the following public GitHub repository: https://github.com/BonellPatinoE/CD48-in-myeloma

## Antibodies

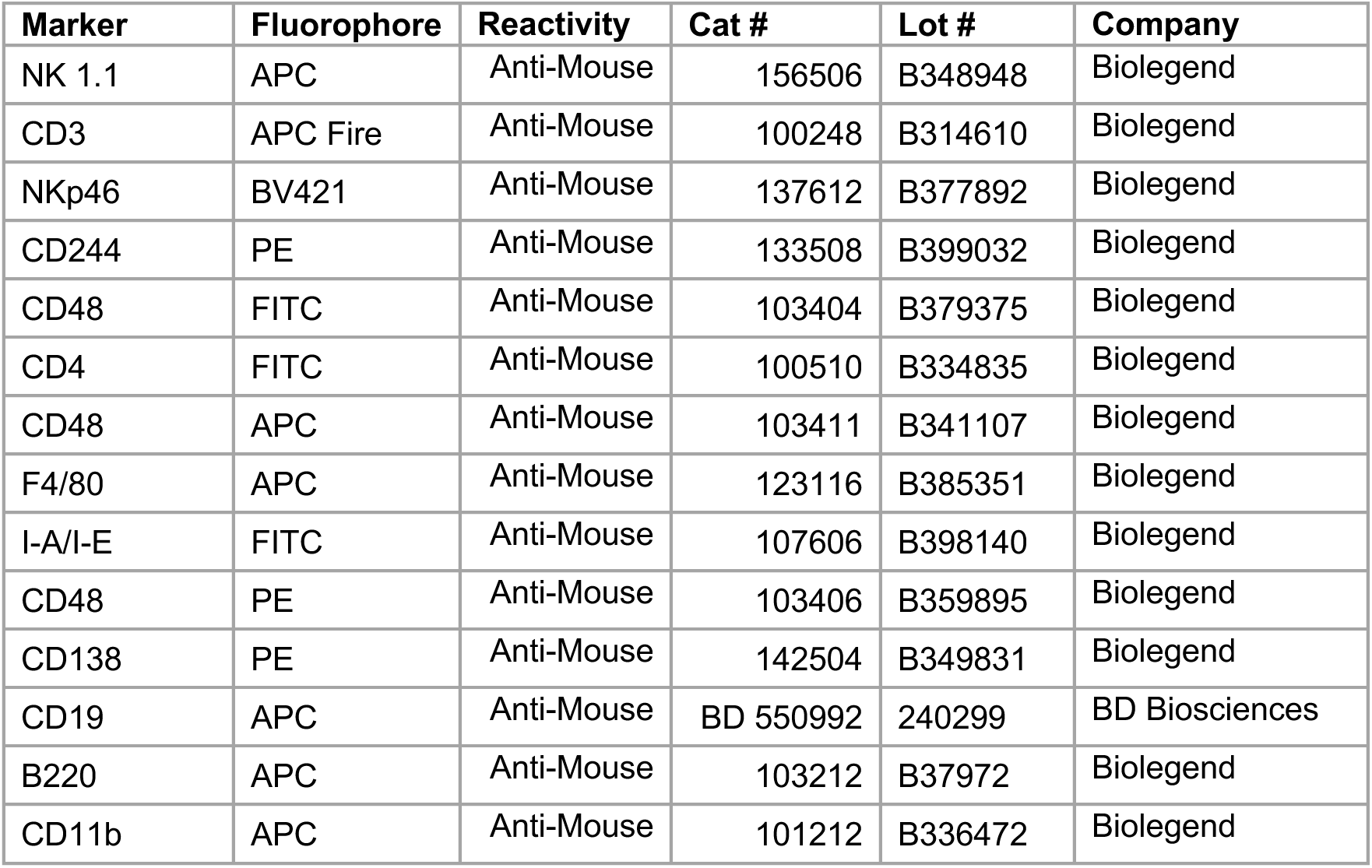
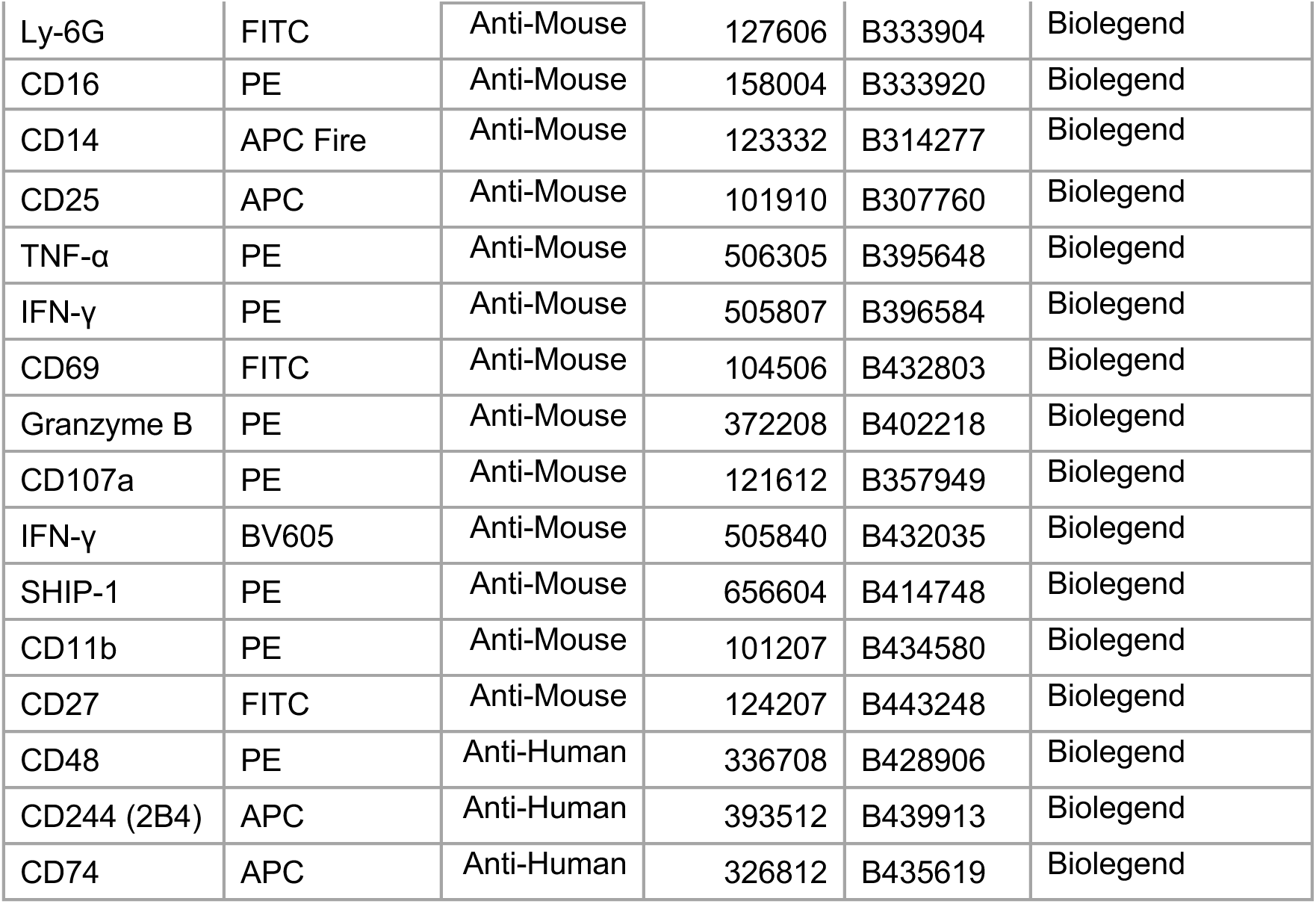

## Purified antibodies and reagents for NK cell Plate-bound assay

| **Antibody** | **Clone** | **Cat#** | **Lot#** | **Company** |
| --- | --- | --- | --- | --- |
| Purified anti-mouse NK1.1 | PK136 | 108702 | B255763 | Biolegend |
| Purified anti-mouse CD335 (Nkp46) | 29A1.4 | 137602 | B382435 | Biolegend |
| Purified anti-mouse CD16 | S17014E | 158002 | B376266 | Biolegend |
| Purified anti-mouse CD244.2 (2B4 B6 alloantigen) | m2B4 | 133501 | B390508 | Biolegend |
| Purified Mouse IgG2a,k isotype Ctrl | MOPC-173 | 400202 | B370572 | Biolegend |
| TrueStain FcX (Anti Mouse CD16/32) | 93 | 101320 | B295040 | Biolegend |
| Brefeldin A |  | 420601 | B406627 | Biolegend |
| Cell Activation Cocktail (without Brefeldin A)<br>[PMA (phorbol 12-myristate-13-acetate) and ionomycin] |  | 423302 |  | Biolegend |
| Monensin |  | 420701 | B431167 | Biolegend |

