## Supplemental figures for "Integrative Genomic, Single-Cell, and Functional Profiling of the CD48–CD244 Axis and NK-Cell Dysfunction in Multiple Myeloma"

### Supplementary Figure S1

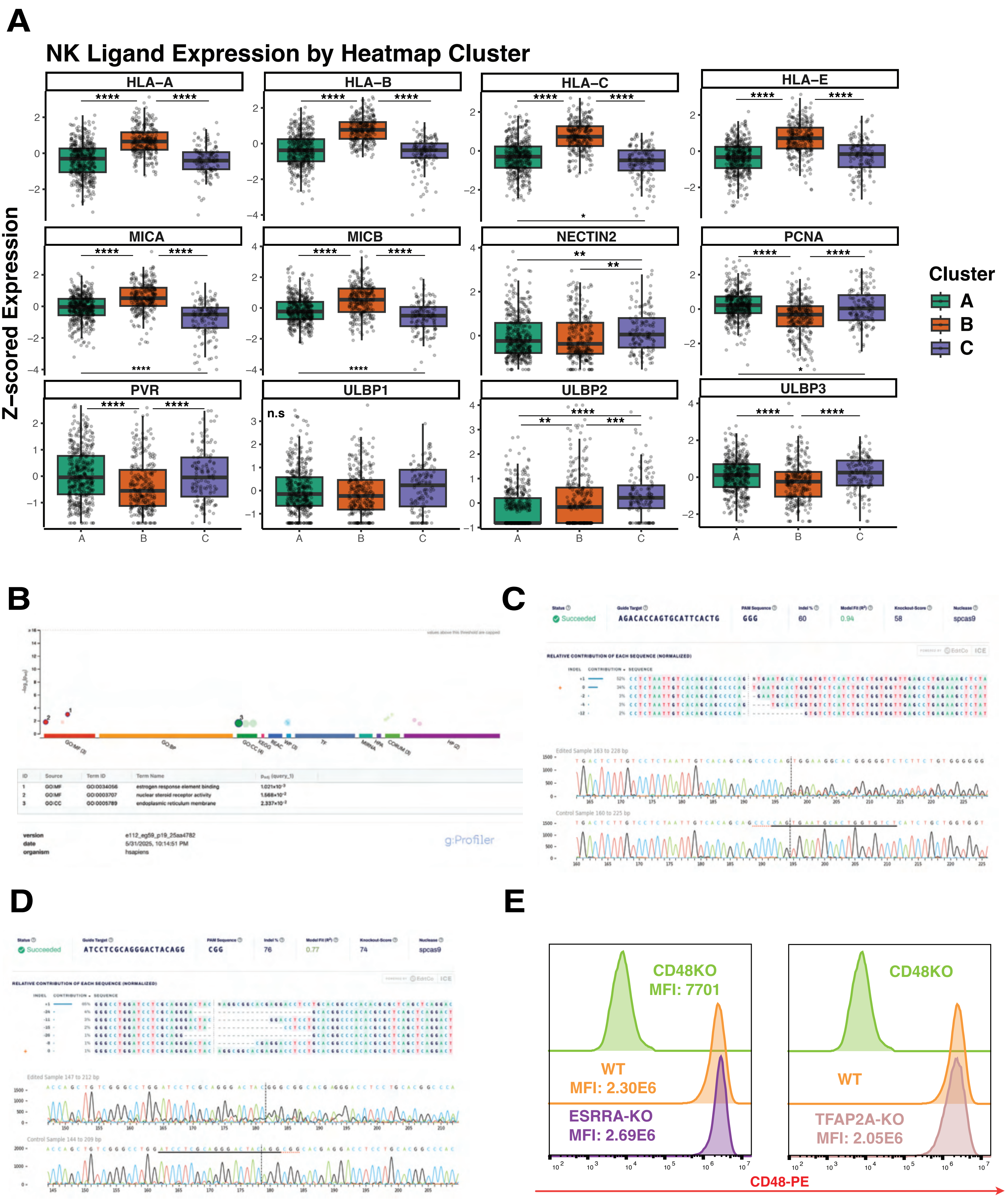

**Supplementary Figure S1. NK Ligand Expression and Validation of CD48 Regulators Identified by CRISPR Screening. A.** Boxplots showing z-scored expression of NK cell ligands in tumor cells from the CoMMpass IA19 dataset across transcriptomic clusters (A, B, C) identified by heatmap-based clustering of myeloma patient samples. Statistical significance was assessed by one-way ANOVA; ****$p < 0.0001$. **B.** Functional enrichment analysis using gProfiler of genes identified in genome-wide CRISPRa/KO screens as regulators of CD48 expression. **C–D.** Validation of CRISPR-mediated knockout of ESRRA (C) and TFAP2A (D) using Synthego ICE analysis and Sanger sequencing, showing indel spectra and editing efficiencies. **E.** Flow cytometry quantification of CD48 surface expression (MFI) in WT, ESRRA-KO, and TFAP2A-KO clones compared to CD48 knockout cells.

### Supplementary Figure S2

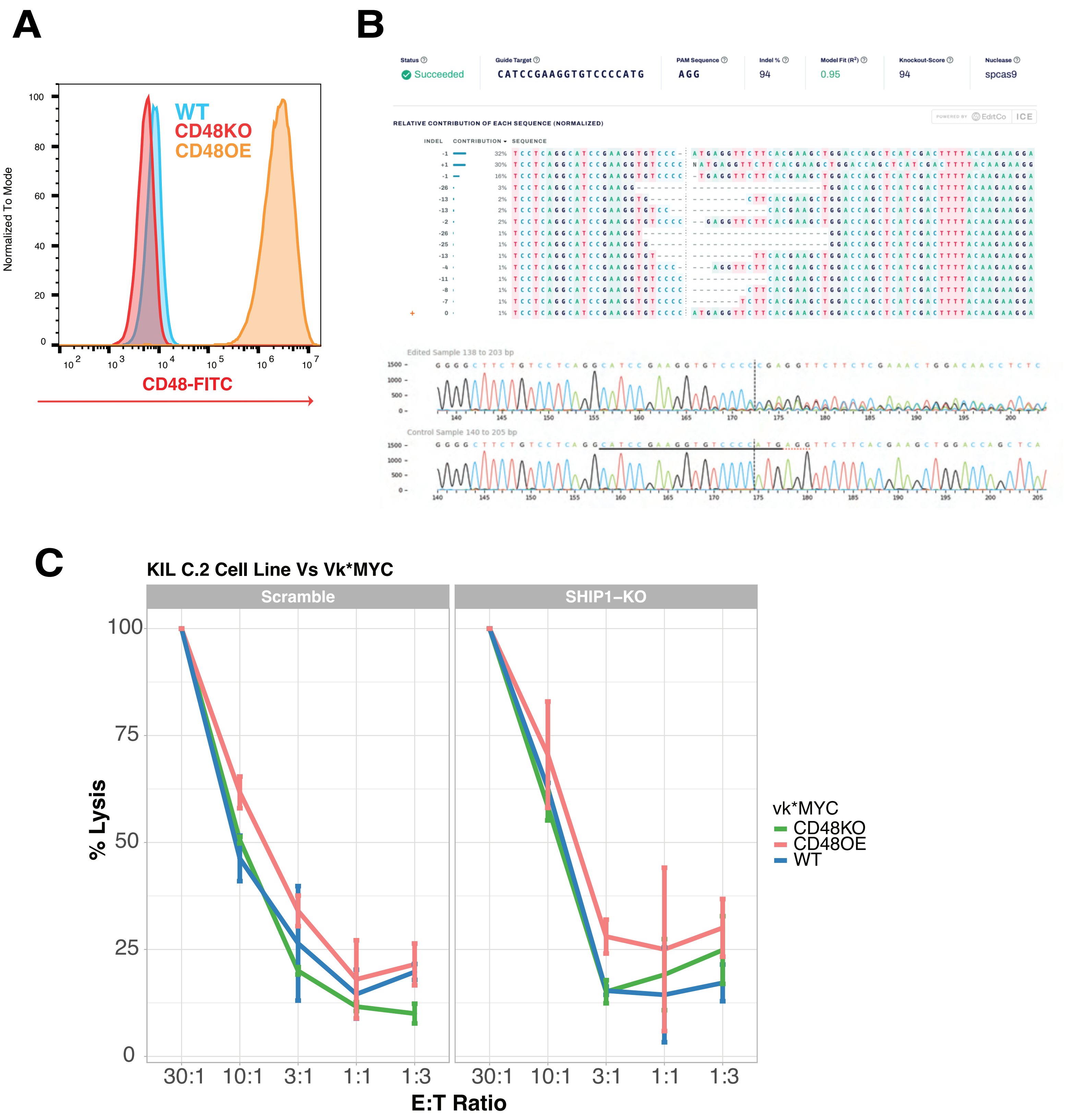

**Supplementary Figure S2. Functional and Genotypic Validation of CD48 Manipulation in Murine vk*MYC Cells. A.** Flow cytometry histogram showing surface CD48 expression in WT, CD48KO, and CD48OE vk*MYC cells. **B.** ICE analysis confirming efficient CRISPR-mediated knockout of INPP5D (SHIP1), showing indel spectrum and Sanger chromatograms. **C.** NK cell cytotoxicity assays using KIL C.2 cells (Scramble or SHIP1-KO) against Vk*MYC targets (WT, CD48KO, CD48OE). Percent lysis is plotted across E:T ratios.

### Supplementary Figure S3

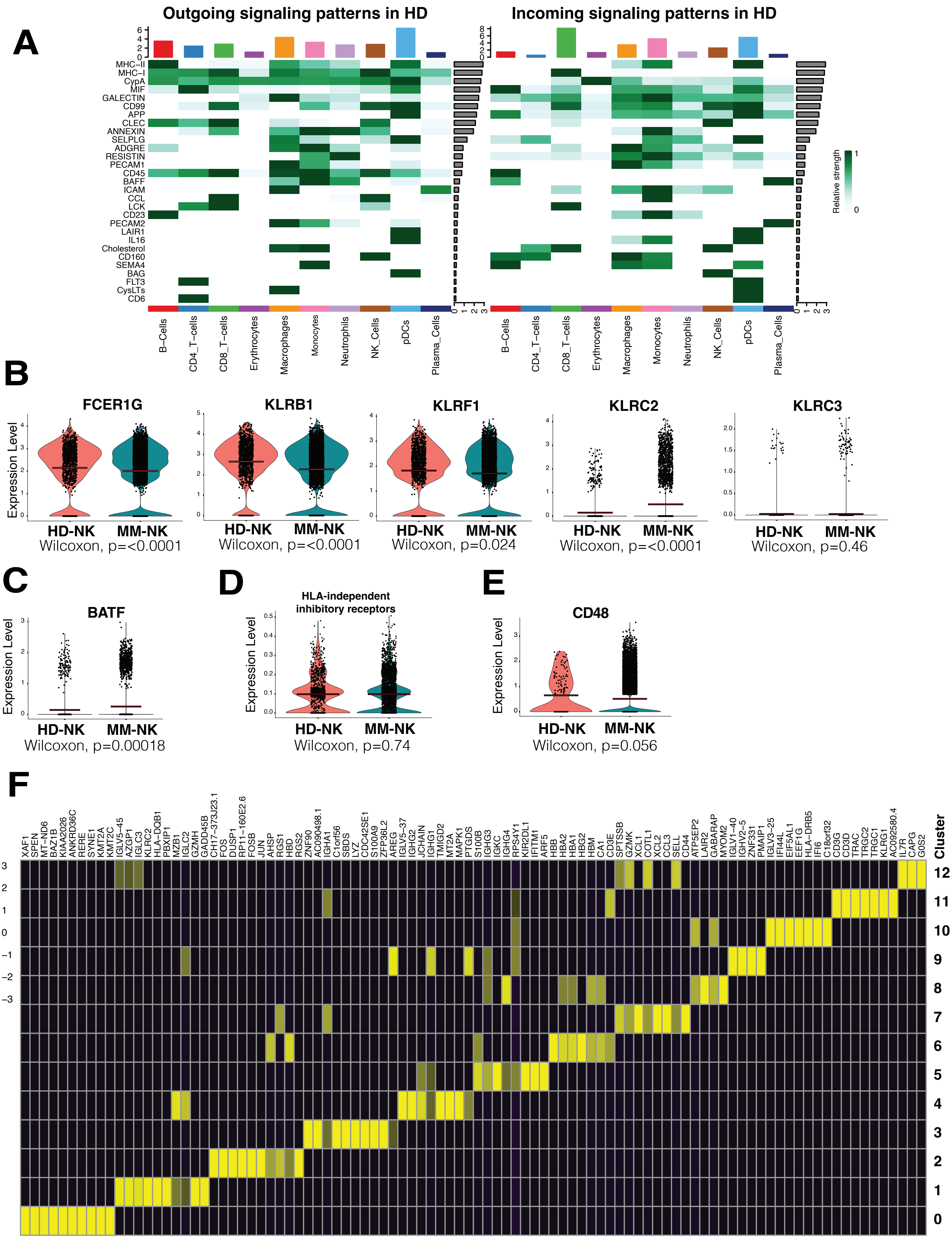

**Supplementary Figure S3. Signaling Landscape and "CMV adaptive-like" CD56–dim phenotype markers Signatures in NK Cells from HD vs MM. A.** CellChat analysis showing outgoing and incoming signaling patterns across immune subsets from healthy donor (HD) bone marrow. **B.** Violin plots comparing expression of "CMV adaptive-like" CD56–dim phenotype markers (FCER1G, KLRB1, KLRF1, KLRC2, KLRC3) in HD-NK vs MM-NK cells; p-values by Wilcoxon test. **C.** BATF expression HD-NK and MM-NK cells. **D.** Violin plot showing no significant difference in HLA-independent inhibitory receptors score between HD-NK and MM-NK cells and **E.** Violin plot showing CD48 expression in HD-NK and MM-NK. **F.** Heatmap displaying cluster-specific marker gene expression across NK-cell subclusters.

### Supplementary Figure S4

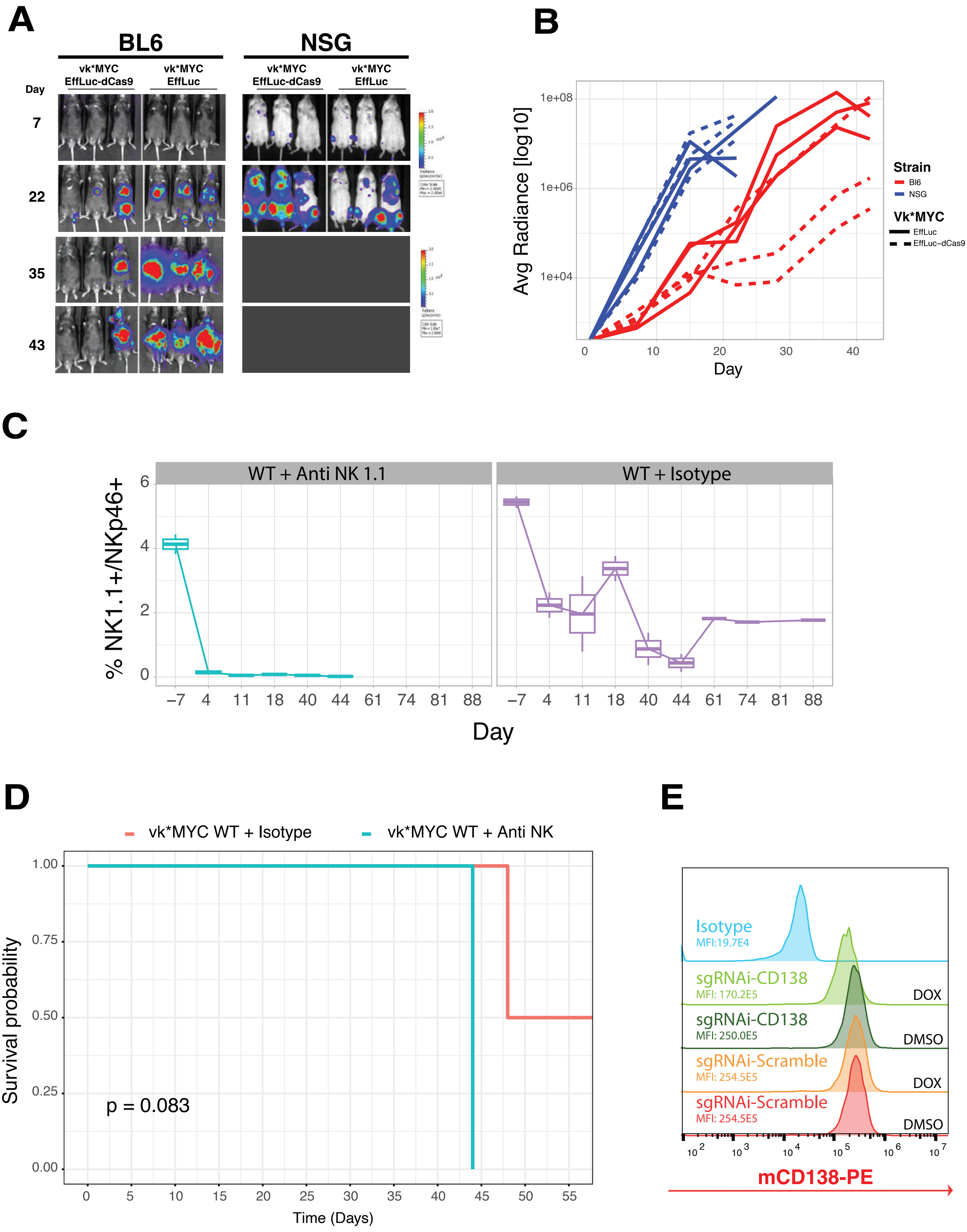

**Supplementary Figure S4. In Vivo Modeling of CD48 Function Using the Vk*MYC Luciferase and dCas9-KRAB System. A.** Representative in vivo bioluminescent images of VkMYC-bearing BL6 and NSG mice. Tumor cells expressing firefly luciferase and/or dCas9-KRAB demonstrate successful engraftment and allow for tumor tracking without the need for serum protein electrophoresis (SPEP), in a non-secretory Vk*MYC clones (Vk32245). **B.** Longitudinal quantification of average tumor radiance (log10 scale) in each condition, grouped by mouse strain and tumor modification. **C.** Flow cytometry quantification of $NK1.1^{+}NKp46^{+}$ NK cells over time, comparing NK cell depletion (anti-NK1.1) versus isotype control in vk*MYC WT. **D.** Kaplan–Meier survival curves comparing Vk*MYC WT mice with NK cell depletion versus isotype control. Statistical significance was assessed using the log-rank test. **E.** Flow cytometry analysis of CD138 expression in Vk*MYC cells engineered with a doxycycline-inducible sgRNA in a dCas9-KRAB constitutive system targeting CD138 (sgRNA-CD138), compared to scramble sgRNA control.

### Supplementary Figure S5

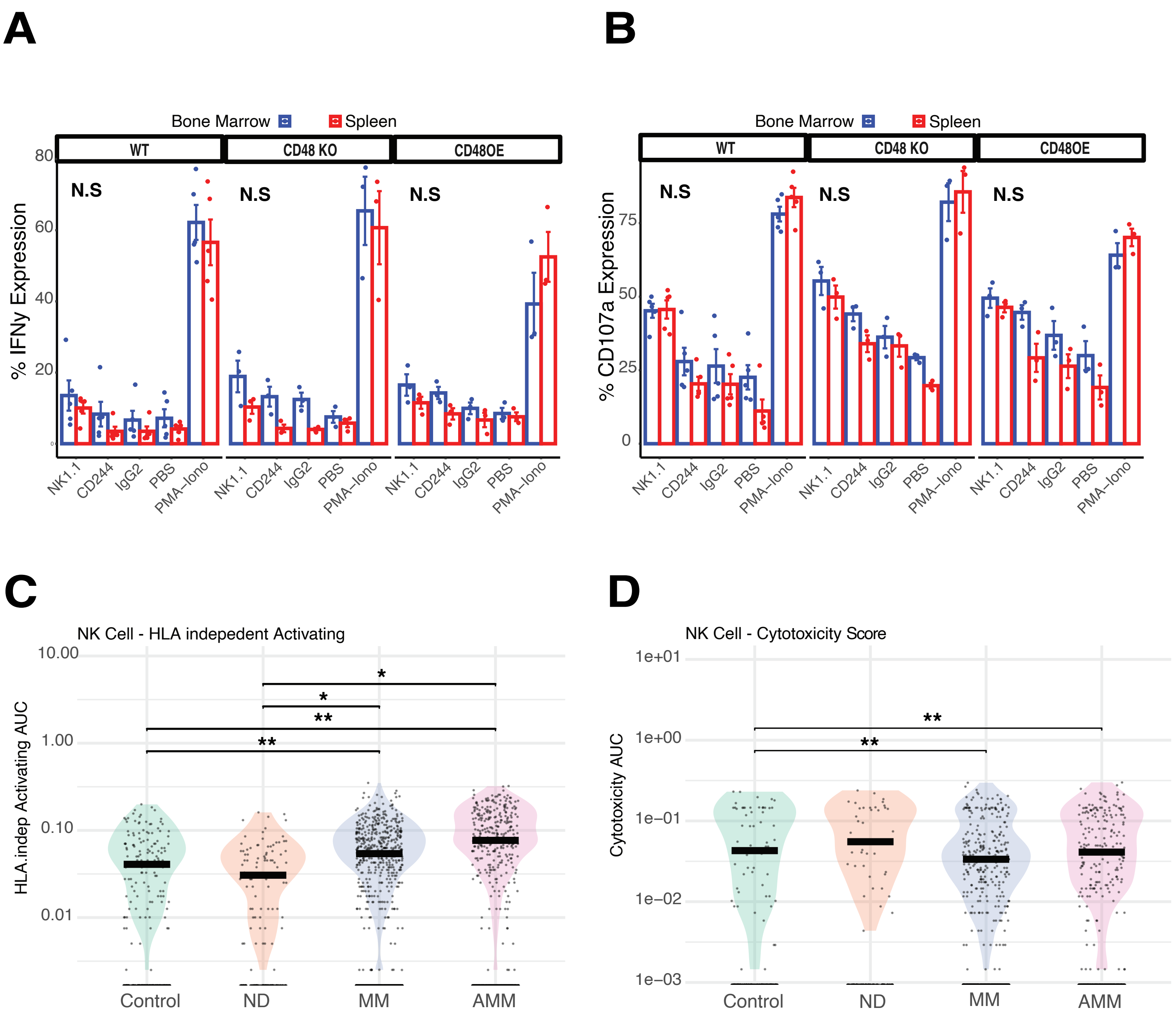

**Supplementary Figure S5. NK Cell Functional Output and Transcriptional Programs in the Vk*MYC Mouse Model. A.** Percentage of IFN-γ–producing NK cells in the bone marrow (blue) and spleen (red) of mice bearing WT, CD48KO, or CD48OE Vk*MYC tumors following ex vivo stimulation with anti-NK1.1, CD244, or PMA/ionomycin. No significant differences (N.S.) were observed across tissues. **B.** Percentage of CD107a$^{+}$ degranulating NK cells under the same conditions as in panel A. CD48 manipulation did not significantly affect NK cell degranulation in either compartment. **C.** AUCell-based HLA-independent NK cell activation scores derived from single-cell RNA-seq data of tumor-infiltrating NK cells in Control, newly diagnosed (ND), multiple myeloma (MM), and advanced MM (AMM) samples from the immunocompetent Vk*MYC model. **D.** AUCell-based NK cell cytotoxicity scores across the same groups as in panel C. Statistical significance was determined using the Wilcoxon rank-sum test; $*p < 0.05$, $**p < 0.01$.
